# Bioenergetic And Protein Processing Imbalances Synergize in iPSC-Dopamine Neurons from Individuals with Idiopathic Parkinson’s Disease

**DOI:** 10.1101/2025.07.08.663742

**Authors:** Kelsey Bernard, Mandi J Corenblum, Paola Tonino, Lalitha Madhavan

## Abstract

Patient induced pluripotent stem cell (iPSC)-based models represent a powerful human system to gain insights into the etiopathology of Parkinson’s disease (PD). Here, we study several iPSC-derived dopamine neuron (iPSC-DAN) lines, from individuals with idiopathic PD, which is the most common form of PD. Specifically, using iPSC-DAN differentiated for 50-55 days, we performed an in-depth analysis of different bioenergetic pathways and cellular quality control mechanisms in the cells. Our results showed wide ranging impairments in oxidative phosphorylation (OXPHOS), glycolysis and creatine kinase pathways in the PD dopamine (DA) neurons. Specifically, the PD neurons exhibited reduced oxygen consumption rates (OCR) at baseline and after challenges with mitochondrial inhibitors, as well as decreased glycolytic reserves measured via ECAR. This translated to lower OCR:ECAR ratios signifying more reliance on glycolysis vs OXPHOS in the PD cells. Moreover, a mislocalization of creatine kinase B to mitochondria was seen in the PD cells. These energetic changes synergized with the enhanced expression of mitochondrial fission proteins, disrupted mitophagy and oxidative stress. Additionally, the PD neurons contained more monomeric, phosphorylated and aggregated forms of alpha synuclein and displayed reduced viability. Ultrastructural examination through immuno-electron microscopy showed more alpha synuclein gold particles directly associated with mitochondria and packing autophagic vesicles. In essence, these data capture a web of key changes in human iPSC-DAN from idiopathic PD subjects associated with neuronal degeneration.

## INTRODUCTION

Parkinson’s disease is the fastest growing neurological disorder and one of the most common progressive degenerative conditions of the nervous system (Bloem *et al*., 2021; Dorsey *et al*., 2018). Although some PD cases are inheritable, the majority of PD develops in a sporadic manner (Corenblum *et al*., 2023; Teves *et al*., 2018). Understanding the etiopathology and molecular basis of neurodegeneration in PD remains a fundamental pursuit; however, the absence of appropriate models has been a major bottleneck in the field. From this viewpoint, human induced pluripotent stem cells (iPSCs) have emerged as an important tool to study PD (Badger *et al*., 2014; Caiazza *et al*., 2020). We recently reported that iPSC-derived dopamine neurons (iPSC-DAN), from individuals with sporadic PD, show different pathological phenotypes (Corenblum *et al*., 2023), thus generating a human cellular system in which to more deeply address sporadic PD mechanisms.

Bioenergetic and protein processing defects are proposed as important triggers in the PD disease process (Johnson *et al*., 2019; Malkus *et al*., 2009). Midbrain dopamine (DA) neurons are metabolically demanding because of their intrinsic pacemaker ability, large axonal field and unique somatodendritic characteristics (Matsuda *et al*., 2009; Surmeier *et al*., 2011). Mitochondria produce most of the cell’s ATP through oxidative phosphorylation (OXPHOS) to meet these energy needs (Mosharov *et al*., 2025; Singh *et al*., 2019; Teves *et al*., 2018). Initially, glucose is converted into pyruvate during glycolysis, which occurs in the cytoplasm, producing a small amount of ATP (Cunnane *et al*., 2020; Requejo-Aguilar and Bolanos, 2016; Song *et al*., 2024). This pyruvate enters mitochondria and is processed through the Krebs cycle to generate high energy intermediates which are then fed into the electron transport chain during oxidative phosphorylation. Reactive oxygen species (ROS) are a byproduct of oxidative phosphorylation (Murphy, 2009; Venditti *et al*., 2013), thus also making mitochondria important players in oxidative stress generation. In addition to glycolysis and oxidative phosphorylation, a third system, namely the creatine phosphocreatine shuttle acts to rapidly replenish ATP by transferring high-energy phosphate groups from phosphocreatine (PCr) to ADP, via the enzyme creatine kinase, to maintain energy homeostasis particularly in areas with greater energy requirements (Wallimann *et al*., 2011). Essentially, the creatine kinase (CK) shuttle serves as an energy transport system to transfer high-energy phosphate groups from mitochondria (where ATP is produced) to areas of more energy demand like the cytoplasm, where ATP is utilized (Jost *et al*., 2002). Furthermore, fission and fusion processes dynamically maintain mitochondrial structure and homeostasis, and are naturally linked to the mitochondria’s energy producing function. However, alterations in these interconnected mitochondrial structure and function aspects of PD are not well studied in the human context.

Bioenergetic compromises also appear to be closely related protein deposition in neurodegenerative scenarios. Neurons are known to be highly dependent on the removal of damaged and obsolete cellular components through autophagy. In fact, a prevalent pathological feature of PD is the compromise of mitochondrial quality through aberrant selective autophagy (mitophagy), and the accumulation of misfolded alpha synuclein (αSyn) which also relates to declines in broader cellular autophagy mechanisms (Evans and Holzbaur, 2020; Fleming *et al*., 2022; Palmer *et al*., 2025; Stavoe and Holzbaur, 2019). Specifically, mitochondrial dysfunction is linked to the impairment of autophagy, through the accumulation of damaged mitochondria because of defective mitochondrial turnover, as well as through the depletion of functional lysosomes which are core to the autophagy process (Fleming *et al*., 2022; Palmer *et al*., 2025). This indicates that the different molecular pathways involved in PD pathogenesis may be ultimately related (Poewe *et al*., 2017).

From this perspective, here, we used multiple human midbrain dopamine (DA) neuron lines, differentiated from iPSCs of idiopathic PD and healthy control subjects, to deeply investigate interrelated bioenergetic and protein quality control processes. Specifically, we examined different energetic systems, mitochondrial dynamics, autophagy, and αSyn biology. Our results reveal distinct alterations in different mitochondrial and protein level processes which seem to dovetail into each other to reduce the survival of PD iPSC-derived DA neurons (iPSC-DAN).

## METHODS

### iPSC lines and Cell Culture

#### iPSC Lines

Human iPSC lines of idiopathic PD (no known gene mutations) and healthy control subjects were mainly obtained from the Parkinson’s Progression Markers Initiative (PPMI) biorepository at Indiana University (supported by the Michael J. Fox Foundation and its corporate sponsors). All iPSC lines originated from PBMCs and had been reprogrammed via non-integrative episomal vectors. 10 control (C1-C10: 6 Male and 4 Female) and 10 PD (PD1-PD10: 6 Male and 4 Female) lines, which were age-matched, were utilized for the study (**Table 1)**. Clinical and demographic information for each line can be found in **Supplementary Table 1.**

**Table 1:** List of iPSC lines.

| Line Moniker | Subject ID | Gender |
| --- | --- | --- |
| C1 | MJFF-3411 | Male |
| C2 | MJFF-3460 | Male |
| C3 | MJFF-3480 | Female |
| C4 | MJFF-3658 | Male |
| C5 | MJFF-3452 | Female |
| C6 | MJFF-3453 | Female |
| C7 | MJFF-3668 | Male |
| C8 | MJFF-3952 | Female |
| C9 | MJFF-3966 | Male |
| C10 | MJFF-4105 | Male |
| PD1 | MJFF-3419 | Female |
| PD2 | MJFF-3960 | Male |
| PD3 | MJFF-3964 | Male |
| PD4 | MJFF-4051 | Female |
| PD5 | MJFF-4055 | Male |
| PD6 | MJFF-3234 | Male |
| PD7 | MJFF-3459 | Male |
| PD8 | MJFF-3473 | Female |
| PD9 | MJFF-3591 | Male |
| PD10 | MJFF-3664 | Female |

#### Cell culture

As described before (Corenblum *et al*., 2023), iPSCs were cultured and differentiated into midbrain dopamine neurons over a 50-55 day period. Briefly, iPSCs were cultured on hESC-Qualified Matrigel Matrix (Corning, Corning NY) in mTeSR Plus media (Stemcell Technologies, Vancouver BC). Colonies were recurrently clump passaged using Gentle Cell Dissociation Reagent (Stemcell Technologies). Midbrain dopamine (DA) neurons were derived using a modified dual SMAD inhibition based floor plate differentiation method (Chambers *et al*., 2009; Corenblum *et al*., 2023; Kriks *et al*., 2011). Briefly, cells were cultured on Poly-L-Ornithine/Fibronectin/Laminin coated plates [Poly-L-Ornithine: mw30,000-70,000; 15μg/ml, Sigma-Aldrich, Fibronectin: 4μg/ml, Corning, Laminin: 6ug/mL, Sigma-Aldrich], and neural patterning of the iPSCs was initiated via inhibitors of the SMAD signaling pathway [LDN 193189 (100nM, Tocris, Minneapolis MN) and SB 431542 (10μM, Tocris)], followed by specification towards a ventral midbrain DA neuron fate using morphogens sonic hedgehog (SHH C25II N-Terminus 100ng/mL, R&D Systems, Minneapolis MN), Purmorphamine (2μM, Tocris), and CHIR 99021 (3μM, Tocris). Neuronal maturation was attained through the addition of ascorbic acid, brain derived neurotrophic factor (BDNF; 20ng/mL, Peprotech, Cranberry NJ), glial cell-line derived neurotrophic factor (GDNF; 20ng/mL, Peprotech), transforming growth factor type β3 (TGF-b3; 1ng/mL, Peprotech), DAPT (10uM, Biogems, Westlake Village CA), L-Ascorbic acid (AA; 200uM, Sigma-Aldrich, St. Louis MO), and cyclic AMP (cAMP; 0.5mM, Sigma-Aldrich). Cells were examined at a dopaminergic progenitor stage (25 days of differentiation) and a more mature DA neuron stage (50-55 days of differentiation), at which point they were also subjected to a variety of assays. All iPSC lines were differentiated in parallel and assessed in at least duplicates for all experiments.

### Immunocytochemistry

iPSC-derived DA neurons were dissociated and plated at 40,000 cells per well on German borosilicate glass coverslips coated with Poly-L-Ornithine (15ug/mL Sigma), Fibronectin (4ug/mL Corning), Laminin (6ug/mL Sigma). At maturation, all cells were fixed for 20 minutes at room temperature (RT) with 4% Paraformaldehyde (PFA; Electron Microscopy Sciences, Hatfield PA). For immunocytochemistry, cells were blocked for 2hrs at room temperature [2% normal goat serum, 1% Bovine Serum Albumin, 0.4% Triton-X-100 in PBS] and incubated in primary antibody overnight at 4°C in a humidity chamber. Primary antibodies were detected in a 2-hour incubation at RT with secondary antibodies coupled to fluorochromes Alexa Fluor 488, 594, 647 (Invitrogen, Waltham MA) or a biotinylated secondary (Vector Laboratories, Newark CA) followed by a Streptavidin-tagged Alexa Fluor, and counterstained with 4’,6’-diamidino-2-phenylindole, dihydrochloride (DAPI, Invitrogen). Control conditions constituted the deletion of the primary antibody or secondary antibody and the inclusion of relevant isotype specific antibodies and sera instead of the omitted antibodies. Primary antibodies used were as follows: TH (1:2000); LMX1A (1:250); FoxA2 (1:200); Tuj1 (1:200); Girk2 (1:1000); Map2 (1:8000); αSyn (1:400); TOM20 (1:400); pSyn (1:2000). More antibody details can be found in the table in the **Supplementary Methods** section.

### Microscopy and Cell counts

Fluorescence analysis was performed using a Zeiss Axio-Imager M2 microscope connected to an Axiocam MRm digital camera and Microlucida software (v2019.1.3, MBF Bioscience). Fluorescence images were obtained using a Zeiss LSM880 Inverted confocal microscope (Zeiss, Jena, Germany). Z sectioning was performed at 1–2μm intervals in order to verify the co-localization of markers. Image extraction and analysis was conducted via the Zen Blue software, or FIJI. TOM20/αSyn images were taken at 40X, with a 1.5X zoom. 3 regions were taken from 2 coverslips each for each subject, resulting in ∼6 total images/subject.

#### Cell counts

For the TH and TH/LMX1A cell counts and Tuj1 and Tuj1/FoxA2 cell counts, 5 random fluorescence images were taken at 63X magnification of cells on each of 3 coverslips (n=15) and the number of positively stained cells was elucidated and expressed as percent of DAPI. For TH and TH/Girk2 cell counts, 3 random fluorescence images were taken at 63X magnification of cells on each of 2 coverslips (n=6) and the number of positively stained cells was enumerated and counted as percent of DAPI.

#### Image Analysis

For TOM20 and αSyn, max projected, channel separated images were analyzed independently for each frame. Images in the TOM20 channel were made binary and a standard threshold was applied across all images. Measurements of the total area of positive staining were obtained. For the αSyn channel a similar process was applied, and then the amount of positive αSyn staining above threshold was normalized to the total area of TOM20 staining for the corresponding image. For total αSyn/frame measurements (**Fig. 6)** lower magnification images were used to analyze a larger field of few, and a similar process was used to obtain % area above threshold measurement. Data are represented as mean ±SEM.

### RT-qPCR

Total RNA extraction was carried out from pelleted cells using the Qiagen RNeasy Mini kit (Qiagen, Germantown MD) according to manufacturer’s instructions. RNA was quantified via NanoDrop 2000 spectrophotometry (ThermoFisher, Waltham MA). 1ug of Total RNA was reverse transcribed using Oligo(dT)12-18 primers, dNTP mix, and SuperScript III Reverse Transcriptase reagents (ThermoFisher) in a 20uL reaction. The resultant cDNA was quantified via NanoDrop and stored at -80C. Quantitative PCR was performed using SsoAdvanced Universal SYBR Green Supermix (Bio-Rad, Hercules CA). Each reaction was performed in duplicate in 96-well pcr plates in a reaction volume of 10uL. Each reaction contained 10ng of cDNA, 0.5uM each forward and reverse primers, and 1X Supermix. The reaction protocol starts with a 3-min initial denaturation step at 95 °C, 40 cycles of 95 °C for 15 s and 60 °C for 60 s. Subsequently, the melting curve was verified by amplification of a single product, which was generated starting at 65 °C and increasing to 99 °C by 1 °C for each 30-s cycle. Relative expression was calculated using the ddCT method and values were normalized to GAPDH housekeeping gene.

Primers used were:

TH- Fwd 5’-CGGGCTTCTCGGACCAGGTGTA-3’ Rev 5’-CTCCTCGGCGGTGTACTCCACA-3’
2) Girk2- Fwd 5’-GATGGGAAACTGTGCCTGAT-3’ Rev 5’-CTCCGAGGTCTGTTTGGATTT-3’
3) Nurr1- Fwd 5’-CGCTTCTCAGAGCTACAGTTAC-3’ Rev 5’-TGGTGAGGTCCATGCTAAAC-3’
4) FoxA2- Fwd 5’-CCGACTGGAGCAGCTACTATG-3’ Rev 5’-TACGTGTTCATGCCGTTCAT-3’
5) Gapdh- Fwd 5’AGGTCGGAGTCAACGGATTT-3’ Rev 5’-ATCTCGCTCCTGGAAGATGG-3’.

### Cell Viability

Cellular viability was determined utilizing a standard MTT [3-(4,5-dimethylthiazol-2-yl)-2,5-diphenyltetrazolium bromide] assay (Invitrogen #M6494; Waltham MA), as described previously (Corenblum *et al*., 2023; Teves *et al*., 2018). iPSC-derived DA neurons were detached and processed into single cell suspensions and plated onto 96-well clear flat-bottom plates coated with Poly-L-Ornithine/Fibronectin/Laminin at 80,000 cells/well (at least 4 replicates per experimental run). Cell negative and assay negative controls were also used. At the time of assay, cells were washed once with PBS and placed in a phenol-free version of the culture media. Experimental wells and cell negative wells were then treated with 12mM MTT (prepared as described in assay manual) at 37°C for 2hrs. Assay negative wells received PBS only and were incubated in the same manner. 85uL of the MTT solution was then removed from each well and 25uL Dimethyl sulfoxide (DMSO; Sigma-Aldrich) added to solubilize the generated formazan crystals. Absorbance was read on a standard microplate reader (BMG FLUOstar Omega, Cary NC) at 540nm.

### Reactive Oxygen Species (ROS) measurements

ROS levels were enumerated using the DCFDA/H2DCFDA Cellular ROS Assay Kit (Abcam #ab113851, Waltham MA), as described previously (Corenblum *et al*., 2023; Teves *et al*., 2018). iPSC-derived DA neurons were detached and processed into single cell suspensions and plated onto 96-well black-walled flat-bottom plates at 80,000 cells/well, and included at least 4 replicates per experimental run. Manufacturer’s instructions were followed. Briefly, culture media was removed and 25uM DCFDA solution was added. Cell negative and assay negative controls were also used. Cells were incubated in the DCFDA solution for 45min at 37°C, after which they were washed in PBS, and fluorescence measured at 485Ex/535Em on a standard plate reader (BMG FLUOstar Omega, Cary NC). Values were normalized to protein expression using a standard Bicinchoninic Acid (BCA) assay.

### Seahorse Mito Stress Test

Mitochondrial function was studied as before using a Seahorse XF Cell Mito Stress Test kit (Agilent, Santa Clara CA) (Corenblum *et al*., 2023; Teves *et al*., 2018). iPSC-derived DA neurons were plated in a Seahorse XF24 V7 PS plate at an optimized seeding density of 90,000 cells per well and consisted of at least 5 replicates per line. Cell negative wells were utilized as assay Blank controls. Cells were plated in standard DA neuron maturation culture media in 5% CO2 and 37°C humidified incubator to allow them to adhere and grow in the plate. At assay time, Seahorse XF DMEM medium enriched with 10mM glucose, 2mM L-glutamine and 1mM sodium pyruvate was equilibrated to 37°C with an adjusted pH of 7.35±0.05. All wells were carefully washed 3 times with the Seahorse XF DMEM medium and then incubated for 1hr in assay media in a 37°C CO2-free incubator before the Mito Stress test was conducted. Successive injections of Oligomycin (Oligo; 1uM), Carbonyl cyanide p-trifluoro-methoxyphenyl hydrazone (FCCP; 1uM), and Rotenone/ Antimycin (R/A; 2uM/2uM), all modulators of respiration, were performed in the Seahorse XF Flux Analyzer. Values were normalized to protein concentration via a standard Bicinchoninic Acid (BCA) assay.

### Seahorse Glycolysis Stress Test

Glycolytic capacity after glucose starvation was probed using a Seahorse XF Glycolysis Stress Test kit (Agilent, Santa Clara CA). Briefly, iPSC-derived DA neurons were plated in a Seahorse XF24 V7 PS plate at an optimized seeding density of 90,000 cells per well and consisted of at least 5 replicates per line. Cell negative wells were utilized as assay Blank controls. Cells were plated in standard DA neuron maturation culture media in 5% CO2 and 37°C humidified incubator to allow them to adhere and grow in the plate. At assay time, Seahorse XF DMEM medium enriched with 2mM L-glutamine was equilibrated to 37°C with an adjusted pH of 7.4. At assay time, culture media was removed and replaced with the Seahorse XF DMEM medium and then incubated for 1hr in assay media in a 37°C CO2-free incubator before the Glycolysis Stress test was conducted. Successive injections of Glucose (Glycolysis Fuel; 10mM), Oligomycin (ATP synthase inhibitor; 1uM), and 2-DG (Competitive Glucose Inhibitor; 50mM), were performed in the Seahorse XF Flux Analyzer. Values were normalized to protein concentration via a standard Bicinchoninic Acid (BCA) assay.

### TMRE Assay

Adherent iPSC-DAN, at 80,000 cells/well in a 96-well plate, were stained using the TMRE-Mitochondrial Membrane Potential Assay Kit (Abcam) per manufacturer’s instructions. Briefly, control and PD iPSC-DAN were treated with 500nM TMRE for 30 mins at 37°C, and consisted of at least 4 replicates per line. Select wells were pre-treated with 400µM CCCP, a decoupling agent, as a positive control for 45 mins before TMRE. Subsequently, after rinses with PBS fluorescence was measured at 549EX/575Em on BMG FLUOstar (Omega, Cary NC) plate reader.

### Protein Isolation and Preparation of Mitochondrial Enriched Fractions

Proteins were extracted from cells using RIPA buffer (50mM Tris–HCl, 150mM NaCl, 0.1% SDS, 0.5% sodium deoxycholate, 1% Triton X-100, pH 7.4) supplemented with a protease inhibitor cocktail (1:100), 1mM sodium fluoride, and 1mM sodium orthovanadate (Sigma-Aldrich, St. Louis, MO, USA).

Mitochondria were isolated using differential centrifugation. Tissues were homogenized in isolation buffer (225mM mannitol, 75mM sucrose, 0.1mM EDTA, 5mM HEPES, pH 7.4) and centrifuged at 800 × g for 10 minutes at 4°C to remove nuclei and debris. The supernatant was subsequently centrifuged at 8,500 × g for 10 minutes at 4°C. The supernatant at this step was considered to be cytosolic and membrane bound material, and the pellet contained mitochondria. This was followed by an additional clean up step in with isolation buffer with EDTA removed and centrifuged at 10,000 x g for 10 minutes at 4°C to pellet the mitochondria. The mitochondrial pellet was resuspended in a minimal volume for further analysis. Protein concentration was determined using a bicinchoninic acid (BCA) assay.

### Immuno-Blotting

Western blotting was carried out on either whole cell lysates (RIPA extraction) or mitochondria enriched fractions. **Protein Concentration** was measured using Bicinchoninic acid (BCA) assay kit (Pierce, Cat #23225) and absorbance was measured on a FLUOstar Omega plate reader (BMG Labtech, Cary, NC). **Gel Electrophoresis:** 7-20μg of sample was denatured in 2X or 4X Laemmli Buffer and separated via electrophoresis using either 10%, or 12% TGX Hand-cast gels (Bio-Rad, Hercules, CA) on a Bio-Rad Tetra cell System at 100V for 110 min. **Transfer:** After running, gels were transferred to 0.4μm nitrocellulose, or PDVF membranes using a BioRad TurboBlot Transfer system. Membranes were blocked in 5% Bovine Serum Albumin (BSA) or 5% non-fat dry milk (NFDM) in 0.1M TBS with 0.1% Tween-20 for 60 min., and then incubated overnight with primary antibodies with constant agitation at 4°C. After washing in TBS-T, membranes were incubated with the appropriate horseradish peroxidase-conjugated secondary antibody for one hour at room temperature. After washing, membranes were visualized with chemiluminescence (Thermo Scientific SuperSignal West Pico PLUS, cat #PI34580). Blots were scanned on an Azure 500 (AZI500-01, Azure Biosystems, Dublin, CA). Optical Density measurements were obtained using Image Studio Lite Software (LI-COR Biosciences). Obtained measurements were normalized to the amount of loading control within the lane.

### Dot Blotting: Separation of Soluble and Insoluble Fractions

Protein was separated into soluble and insoluble fraction as previously described (Anandhan *et al*., 2021). Cells were subjected to the RIPA buffer described above, and allowed to incubate for 30 min on ice, followed by centrifugation (15,000×*g*, 60 min, 4°C). The supernatant was designated as the *Triton X-100 soluble* fraction. The remaining pellet was again treated with the above RIPA solution, with the addition of 2% SDS, and this fraction was designated as the *Triton X-100 insoluble* fraction. The protein concentration in the different lysates was determined via standard BCA. **Blotting:** 15ug of each lysate was blotted directly onto 0.4um nitrocellulose membrane pre-wetted with TBS and allowed to fully dry for approximately 1 hour. Afterwards, membranes were again wetted in TBS and then incubated in primary and secondary antibodies. Resulting signals were detected with chemiluminescence as described above. **Densitometric analysis:** Chemiluminescent images were imported into FIJI, and circles were hand-drawn around each dot to measure the total intensity of the sample. A measurement of the background intensity was taken and subtracted from each sample value. Data is represented as the mean ± SEM.

### Immuno-Electron Microscopy

Immunogold labeling of αSyn was conducted according to the pre-embedding technique with modifications. In brief, day 52 control and PD iPSC-DAN lines were rinsed with 10 mM Phosphate Buffer Saline (PBS), pH 7.2, at room temperature (RT), scraped off well bottoms, transferred to 1.5mL Eppendorf tubes. After centrifugation at 1000 x g, cell pellets were fixed with 4% paraformaldehyde and 0.2% glutaraldehyde in 10mM PBS for 1 hour at RT, washed 3 times with the buffer and followed by aldehyde quenching with 50 mM glycine in 10mM PBS for 15 min. Then, the cell pellets were blocked with a solution of 1% Bovine Serum Albumin (BSA, Jackson ImmunoResearch 001-000-161) in PBS (1% BSA/PBS) containing 0.1% Triton X-100 for 30 min, washed in 1% BSA/PBS for 10 min to remove the Triton X-100 and incubated with monoclonal anti-αSyn antibodies (1:20; BD Biosciences) for 24 hours at 4°C. Negative controls were performed by replacing the primary antibody with 1% BSA/PBS. After primary antibody incubation, the cell pellets were rinsed with the buffer 3 times and incubated with nanogold (1.4nm)-Fab’ goat anti-mouse antibodies (1:40; Nanoprobes, 2002) overnight at 4°C. Next, the cell pellets were rinsed with PBS and gold enhanced for 1 min using GoldEnhance™ EM Plus (Nanoprobes, 2114), according to manufacturer’s instructions. The antibody treated cell pellets were fixed with 3% glutaraldehyde in 10mM PBS for 30 min, rinsed in the buffer and post-fixed in 1% osmium tetroxide in 10mM PBS for 30 min, followed by the processing for routine TEM using Spurr embedding resin. Ultrathin sections (80nm) of cells were obtained in a PTXL ultramicrotome (RMC Boeckeler Instruments, Inc) and contrasted with 2% uranyl acetate and 0.25% lead citrate. Digital images (1792 x 1792 pixels) were acquired in a FEI Tecnai G2 Spirit BT TEM (FEI, Hillsboro, OR) with a side-mounted AMT Image Capture Engine V6.02 (4Mpix) CCD camera, operated at 100 kV.

Quantitative analysis of αSyn immunogold particles was performed on 4 PD and 4 control lines using five images per line. Images were first calibrated using ImageJ software (NIH, Bethesda, MD, USA) after which gold particles number per area (nm^2^) was determined within mitochondria and autophagic vacuoles, or just total gold particles, adjusting image contrast enhancement and segmentation of ROIs by setting a density threshold. Values were expressed as the mean ± SEM.

### Statistical Analysis

GraphPad Prism 10 software (San Diego, CA) was used for statistical analyses. For comparing the two groups, PD vs Control, unpaired *t* tests were applied. In all cases, differences were accepted as significant at *p* < 0.05. The outlier test for The exact sample size and specific statistical details of each experiment of the study are also provided within the relevant result and legend sections.

## RESULTS

### Idiopathic iPSC-DAN neurons show reduced oxidative phosphorylation capacity, reduced viability and dopaminergic differentiation

iPSC lines from persons with idiopathic Parkinson’s disease (PD) and age/sex-matched healthy control individuals (CON) were differentiated into midbrain DA neurons (**Fig. 1A**) and assessed via different output assays between day 50-55. Details of the cell lines and the associated clinical information are provided in **Supplementary Table 1**. A9 subtype ventral midbrain DA neurons were derived from the iPSCs utilizing a modified dual SMAD inhibition floor plate-based approach we have described before (Corenblum *et al*., 2023). Cellular viability of Day 50 iPSC-DAN was elucidated using a MTT assay. A9 Midbrain DA neurons in the lateral tier of the substantia nigra (SN) are known to be particularly vulnerable to degeneration in PD (Barker *et al*., 2015). Similar to our previous study using the same model system, PD neurons showed a significantly reduced viability compared to CON neurons (**Fig. 1B**; p = 0.0004; Unpaired t-test, t = 4.357, df = 18). To examine the bioenergetic profile of the DA neurons, we first applied the Seahorse Mito Stress test (**Fig. 1C**), to analyze mitochondrial respiratory function in both CON and PD iPSC-DAN. Neuronal energy demand is primarily met by mitochondrial production of ATP via oxidative phosphorylation (OXPHOS). The Seahorse Mito Stress test directly measures the oxygen consumption rate (OCR) of cells in the presence of modulators of respiration, specifically Oligomycin, FCCP, Antimycin and Rotenone (Corenblum *et al*., 2023; Gu *et al*., 2021; Teves *et al*., 2018). We found that PD DA neurons had significantly decreased respiration rates at baseline (**Fig. 1D**; p = 0.001; Unpaired t-test, t = 3.926, df =18) and reduced maximal respiration capabilities compared to CON lines (**Fig. 1E**; p = 0.0002; Unpaired t-test, t=4.681, df=18). In conjunction with these OCR data, it was noted that both proton leak (PL) and ATP Production were significantly diminished in PD DA neurons relative to CON neurons (**Fig. 1G** p = 0.0011; Unpaired t-test, t = 3.890, df =18; **Fig. 1F**; p = 0.0026; Unpaired t-test, t = 3.493, df =18, respectively). Spare Respiratory Capacity, which reflects the ‘fitness’ of the neurons by testing their ability to upregulate ATP production in situations of stress or high demand, was also pointedly less in the PD DA neurons (**Fig. 1I**; p = 0.0047; Unpaired t-test, t = 3.221, df = 18). Moreover, the Non-Mitochondrial OCR of PD cells was also lower than CON cells (**Fig. 1H**; p = 0.0007; Unpaired t-test, t = 4.068, df = 18). Overall, these data indicated extensive declines in oxidative phosphorylation capacities in the PD DA neurons.

**Figure 1:**
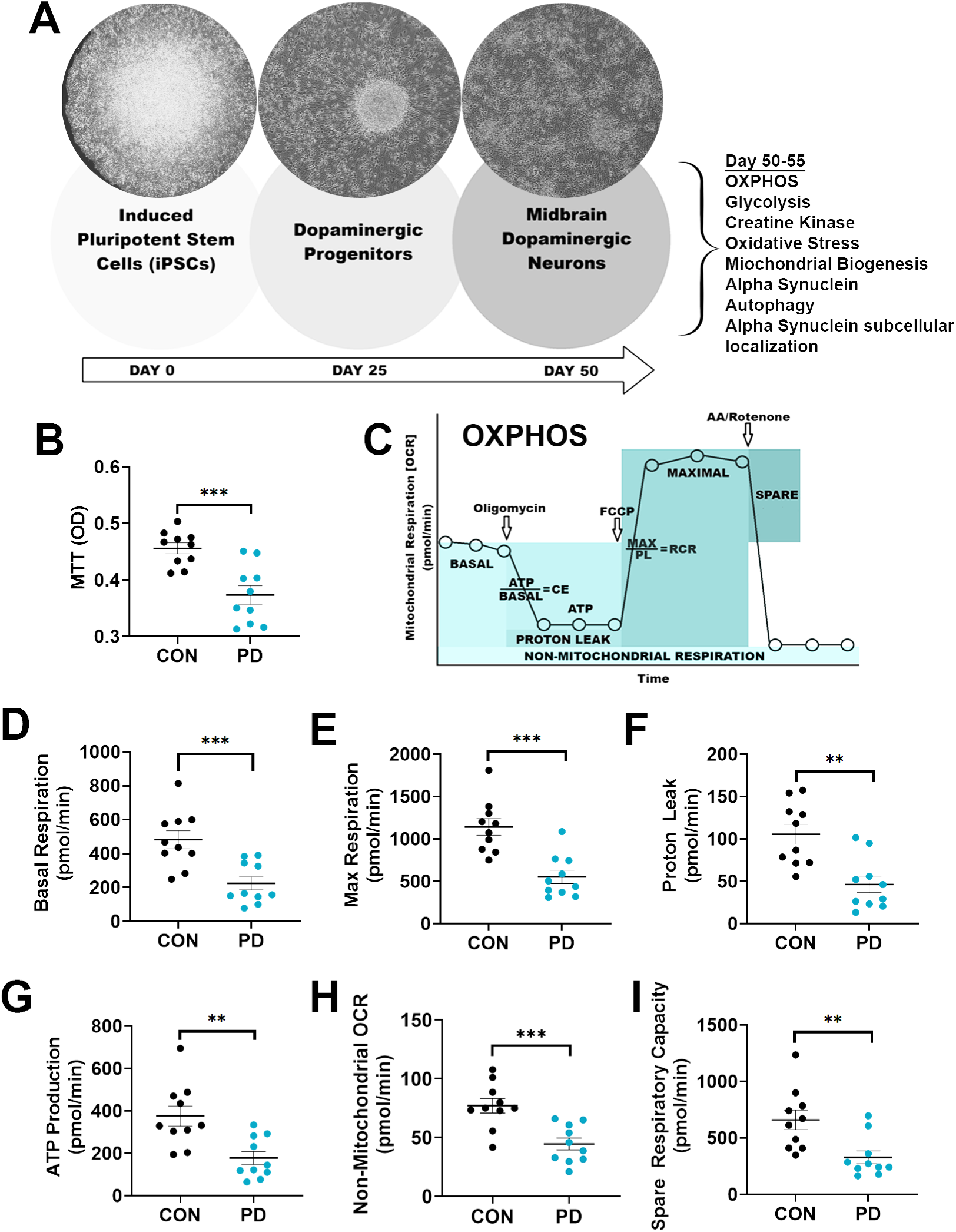
Viability and oxidative phosphorylation assessments in the iPSC-DAN. (A) Schematic representation of experimental timeline and associated assays. Results from a MTT assay comparing the viability of CON vs PD iPSC-DAN is in (B). Schema illustrating parameters elucidated from a Seahorse Mito Stress test is in (C). (D-I) Results obtained on Day 50 iPSC-DAN from the Seahorse Mito Stress test, specifically basal respiration (D), maximal respiration (E), proton leak (F), ATP production (G), non-mitochondrial respiration (H), and spare respiratory capacity (I). *P<0.05, **P<0.01, ***P<0.001, ****P<0.0001, Unpaired t-test; Mean ± SEM, N=10 lines/group

Morphologically, we also saw deficits in the differentiation of the PD DA neurons (**Supplementary Fig. 1**). At day 25 (**Supp Fig. 1A-H**), it was seen that while CON and PD lines differentiated similarly into Tuj1+ neurons (**Supp Fig. 1I**; p = 0.0553; Unpaired t-test, t = 2.049, df =18), the percent of PD cells that co-expressed Tuj1^+^ and LMX1A^+^ (a crucial factor specifying midbrain neuron identity) was significantly less than in the CON lines (**Supp Fig.1J**; p = 0.0002; Unpaired t-test, t = 4.603, df = 18). In addition, it was found that fewer cells from the PD cultures expressed the dopaminergic markers, Tyrosine Hydroxylase (TH) and FoxA2 at this stage (**Supp Fig.1K-T**; TH^+^: p = 0.0108 t = 2.841, df =18, TH^+^/FoxA2^+^: p = 0.0028; Unpaired t-test, t = 3.457, df =18). At 50 days post-differentiation, we examined CON and PD lines for the expression of TH and Girk2. TH^+^/Girk2^+^ co-expression specifically marks the mature A9 subtype of DA neurons located in the lateral tier of the substantia nigra (SN). Our results showed that while there was robust differentiation into A9 DA neurons (**Supp Fig. 1U-b**), there was a significantly lower percentage of TH^+^ and TH^+^/Girk2^+^ co-labeled cells in PD in comparison to control cultures (**Supp Fig. 1c-d**; TH^+^: p = 0.0373, t = 2.271, df =18, TH^+^/Girk2^+^: p = 0.0413; Unpaired t-tests t= 2.220, df = 18). qRT-PCR data supported these results and generally showed lower TH, Girk2, FoxA2 and another important dopaminergic factor, Nurr1 (**Supp Fig. 1e-f).**

### More reliance on glycolysis and lowered glycolytic reserve is seen in the PD DA neurons

We compared OCR to ECAR values from the Mito Stress assay, which measures the rate at which cells release protons (H+) into the surrounding medium and often used as a proxy for glycolytic activity. Cellular reliance on OXPHOS versus Glycolysis can be estimated using such OCR:ECAR ratios. In accordance, at baseline, PD DA neurons showed a significantly decreased OCR:ECAR ratio (**Fig. 2A**; p = 0.0226; Unpaired t-test, t = 2.493, df =18). This originated from the reduced OCR (**Supp Fig. 2A**; p = 0.0014; Unpaired t-test, t = 3.784, df =18) and ECAR (**Supp Fig. 2B**; p = 0.0520; Unpaired t-test, t = 2.081, df =18) seen at baseline. The OCR:ECAR ratio at maximal respiratory capacity was also significantly lower in PD DA neurons (**Fig. 2B**; p = 0.0063; Unpaired t-test, t = 3.094, df =18). Aligning with this, the maximum capacity OCR for PD neurons was substantially less than CON cells (**Supp Fig. 2C**; p = 0.0008; Unpaired t-test, t = 4.037, df =18, while the ECAR maximum showed a reducing trend (**Supp Fig. 2D**; p = 0.1313; Unpaired t-test, t = 1.581, df =18). When post oligomycin OCR:ECAR ratios were computed, it was seen that it was lower in the PD iPSC-DAN, although not statistically significant, compared to CON neurons (**Fig. 2C**; p = 0.1413; Unpaired t-test, t = 1.538, df =18). Correlatively, OCR was found to be reduced in PD DA neurons (**Supp Fig. 2E**; p = 0.0034; Unpaired t-test, t = 3.366, df =18), as was the ECAR (**Supp Fig. 2F**; p = 0.0608; Unpaired t-test, t = 2.000, df =18). These OCR:ECAR ratios suggested that the PD iPSC-DAN maybe more reliant on glycolysis than CON cells.

**Figure 2:**
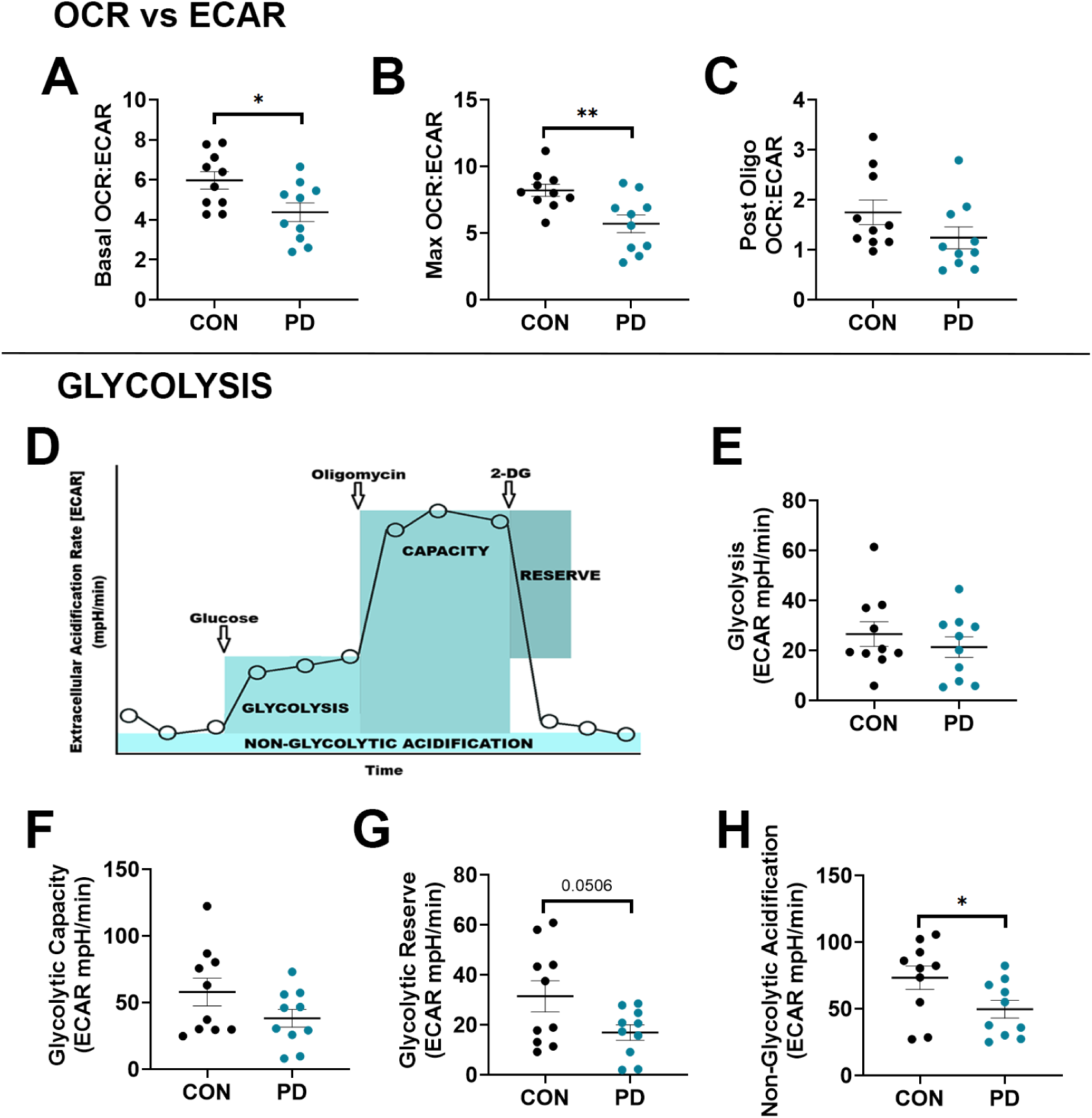
OCR vs ECAR and Glycolysis measurements. Neuronal reliance on oxidative phosphorylation versus glycolysis was analyzed through OCR:ECAR ratios. OCR to ECAR ratios computed at Baseline (A), at Maximal capacity (B), and post Oligomycin administration (C). (D) depicts a schematic representation of Glycolysis Stress test outcome measures. (E-H) show ECAR results from the Glycolysis Stress test for overall Glycolysis (E), Glycolytic Capacity (F), Glycolytic Reserve (G), and Non-Glycolytic Acidification (H) between control and PD iPSC-DAN. *P<0.05, **P<0.01, ***P<0.001, ****P<0.0001, Unpaired t-test; Mean ±, SEM N=10 lines/group.

To investigate the above results further, we performed a Glycolysis Stress test to assess key parameters of glycolytic flux by directly measuring the extracellular acidification rate (ECAR) in cells (**Fig. 2D**). After a saturating injection of glucose, CON and PD DA neurons had similar ECAR values (**Fig. 2E**; p = 0.4312; Unpaired t-test, t = 0.8052, df =18), suggesting that the rate of glycolysis under basal conditions was not significantly altered. Subsequent injection of the ATP synthase inhibitor, Oligomycin, revealed a slight decrease in the cellular maximum glycolytic capacity of the PD DA neurons compared to the CON cells (**Fig. 2F**; p = 0.1279; Unpaired t-test, t = 1.596, df =18). With the addition of 2-deoxy-glucose (2-DG), a glucose analog, the calculated difference between the glycolytic capacity and rate of glycolysis showed that the glycolytic reserve in PD DA neurons was notably less than in CON lines (**Fig. 2G**; p = 0.0506; Unpaired t-test, t = 2.095, df =18). Interestingly, non-glycolytic acidification, due to other metabolic processes than glycolysis, was also found to be significantly decreased in PD neurons (**Fig. 2H**; p = 0.0459; Unpaired t-test, t = 2.145, df =18).

### Creatine kinase energetics are less efficient in the PD iPSC-DAN

Neurons are creatine containing cells. The Creatine Kinase (CK) energy shuttle connects sites of ATP production (glycolysis and mitochondrial oxidative phosphorylation) with subcellular sites of ATP utilization (ATPases) (Jost *et al*., 2002; Wallimann *et al*., 2011). This involves the reversible transfer of a phosphate group between ATP and creatine, forming phosphocreatine (PCr) and ADP. Thus, diffusion limitations of ADP and ATP are overcome by this shuttle to quickly make ATP available where needed in the cell. CK is expressed in the cytosol, as well as in the mitochondria of cells. (**Fig. 3A**). There is a slight, but not significant (p = 0.0769, Unpaired t-test, t = 1.883, df =17) reduction in CKB in whole cell lysates of PD lines (0.883 ± 0.179), relative to CON (1± 0.121) (**Fig. 3B-C**). To look specifically within the mitochondrial compartment, cells underwent enrichment for the mitochondrial fraction through a sucrose extraction process (**Fig. 3D**). The resulting lysates were probed for CKB as well as mitochondrial specific uMt-CK. There was no change to the amount of uMt-CK (**Fig. 3E-F**), but surprisingly, we noted a significant increase in the CKB localized to this region (**Fig. 3E-G**; CON = 0.9773± 0.4829 vs. PD = 1.711± 0.8062, p = 0.356, Unpaired t-test, t = 2.309, df =15).

**Figure 3:**
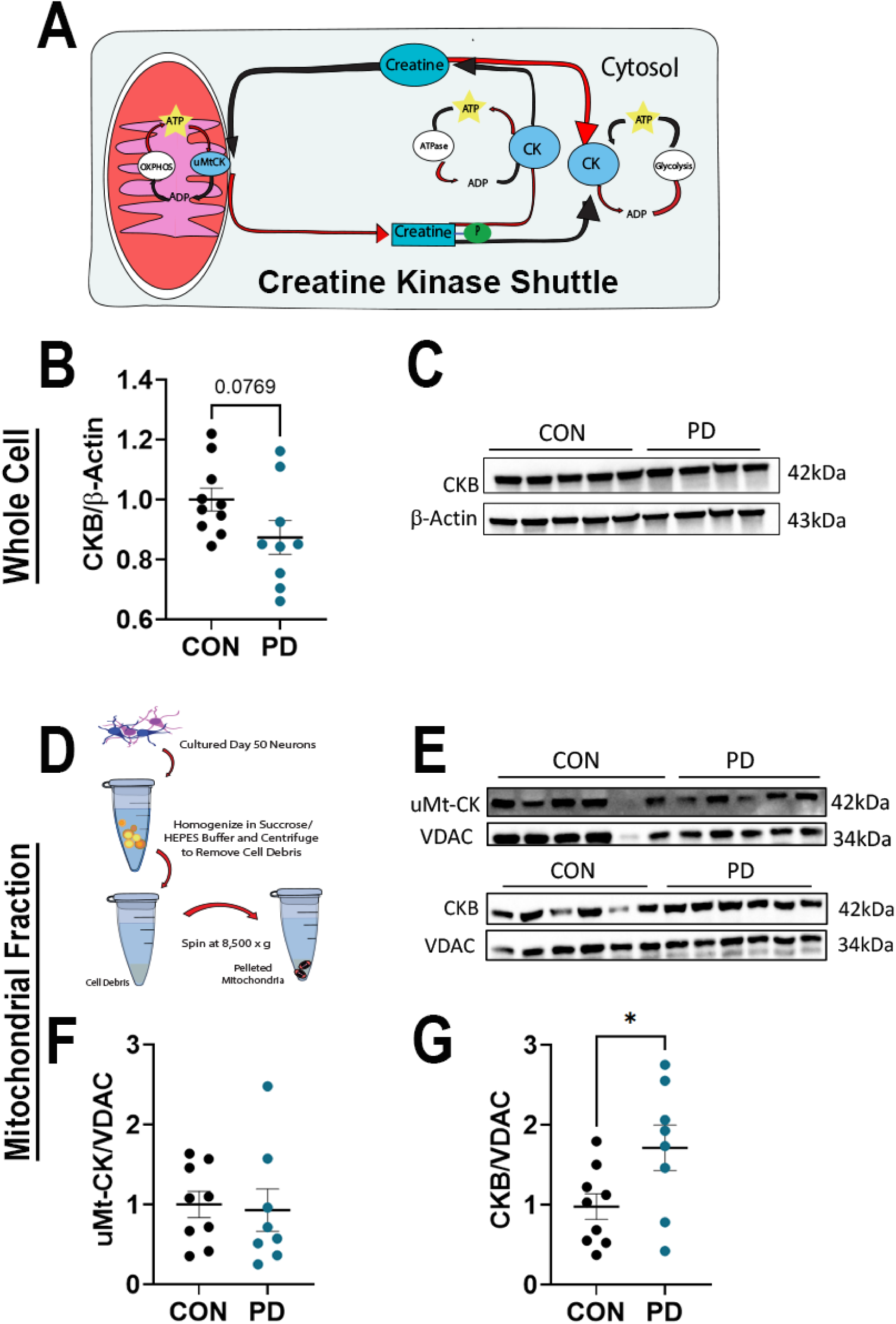
Creatine kinase expression in the iPSC-DAN. Schematic in (A) outlines the creatine kinase shuttle. (B) Quantification of CKB protein analysis via western blotting of whole cell lysates. Representative blots from CON and PD DAN are shown in (C). Mitochondrial enrichment procedure is schematized in (D), with corresponding western blots shown in (E) for uMt-CK and CKB. Associated quantifications are in (F-G). *P<0.05, **P<0.01, ***P<0.001, ****P<0.0001, Unpaired t-test; Mean ± SEM, N-8-10 lines/group.

### Cellular redox balance is altered in the PD DA neurons

Changes to mitochondrial functioning and ATP production can result in excess reactive oxygens species (ROS). Given the alterations of mitochondrial ATP production in PD iPSC-DAN relative to CON (**Figures 1 & 2**), we next assessed the presence of ROS. We used DCFH-DA, a fluorescent probe that is oxidized in the presence of peroxide species (**Fig. 4A**). PD iPSC-DAN displayed increased fluorescent intensity (**Fig. 4B**; PD = 6833 ± 1107 au, vs CON = 5581 ± 493.2 au; t = 3.246, df = 17, p = 0.0048, Unpaired t-test). Excess ROS can damage lipid membranes, alter DNA, and damage cellular proteins. Given this, cells are equipped with several antioxidant response mechanisms that are upregulated after ROS detection. We probed NAD(P)H quinone dehydrogenase-1 (NQO-1), glutathione-S-transferase mu-1 (GSTM-1), glutamate-cystine synthetase (γGCSm), and heme-oxygenase-1 (HO-1), to examine antioxidant changes (**Fig. 4C-J)**. There was no difference in the amount of NQO-1 in CON (0.648±0.7) compared to PD cells (0.526 ± .327) (**Fig. 4C, E**). Similarly, there were no changes in the amount of γGCSm (**Fig. 4D, F**) or HO-1 (**Fig. 4H, J**). However, there was a marked reduction in the amount of GSTM-1 (**Fig. 4G, I**; CON = 1.0±0.8191 vs. PD = 0.2167 ± 0.1181, t=2.66, df = 16). In summary these data show increased ROS but lesser antioxidant capacity in the PD DA neurons.

**Figure 4:**
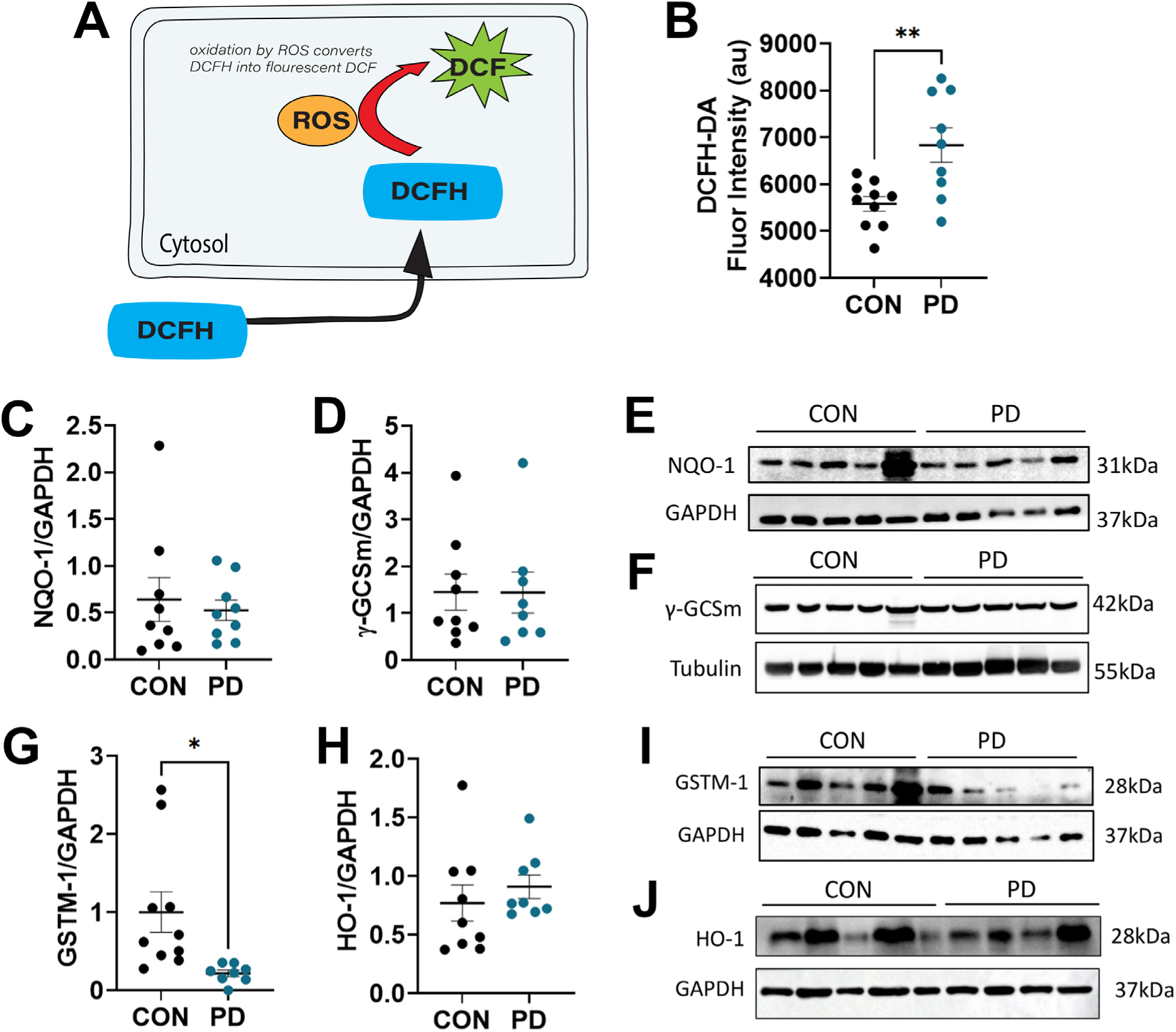
Reactive oxygen species and antioxidant expression. Schematic depiction of the DCFH-DA assay (A). ROS quantification of Day 50 DAN from CON and PD cells (B). Western blot results from antioxidant molecules NQO-1 (C), γGCSm (D), GSTM-1 (E), and HO-1 (F). Representative blots are shown in (E-F, I-J) respectively. *P<0.05, **P<0.01, ***P<0.001, ****P<0.0001, Unpaired t-test; Mean ± SEM, N=8-10/lines/group

### PD DA neurons show dysfunctional mitophagy and mitochondrial dynamics

To better understand the changes to the energy metabolism noted, we assessed the structural competence of the mitochondrial membrane, as well as several proteins involved in mitochondrial dynamics. Mitochondria display a steep ionic gradient across their membrane, necessary for the production of ATP (Vercellino and Sazanov, 2022; Zorova *et al*., 2018). Several proteins, including Pink1 and Parkin act as a part of quality control of these organelles by sampling the charge across the membrane, and targeting the mitochondria for degradation with insufficient potential (**Fig. 5A**) (Pickrell and Youle, 2015). Tetramethylrhodamine ethyl ester (TMRE) is a positively charged fluorescent molecule that accumulates in the inside of negatively charged mitochondria. Loss of fluorescence indicates a lack of negative charge within mitochondria, an indicator of collapsed potential across the membrane. PD iPSC-DAN subjected to treatment with TMRE showed a reduction in overall fluorescence (**Fig. 5B**; t=3.390, df=17; p=0.035, Unpaired t-test). Additionally, whole cell lysates underwent electrophoresis, were immuno-blotted for Pink1 and Parkin (**Fig. 5C-E**), and densitometric values normalized to the amount of GAPDH within the lane. As seen, there is an overall reduction in the level of Pink1 detected (**Fig. 5C**) (CON = 1.0± 0.491, PD = 0.5355 ± 0.3392, t = 2.312, df = 16, p = 0.0340). Conversely, there was a noted increase in Parkin found in these cells (**Fig. 5D**) (CON = 1.0 ± 0.3339, PD = 3.315 ± 2.724, t = 2.674, df =17, p = 0.016). To probe the sub-cellular localization of these proteins, mitochondria were enriched for using the previously described method and subjected to immuno-blotting. Interestingly, there was an increase in Pink1, and a decrease in parkin (both not statistically significant) relative to the mitochondrial specific house-keeping protein VDAC within this fraction (**Fig. 5F, G, H**).

**Figure 5:**
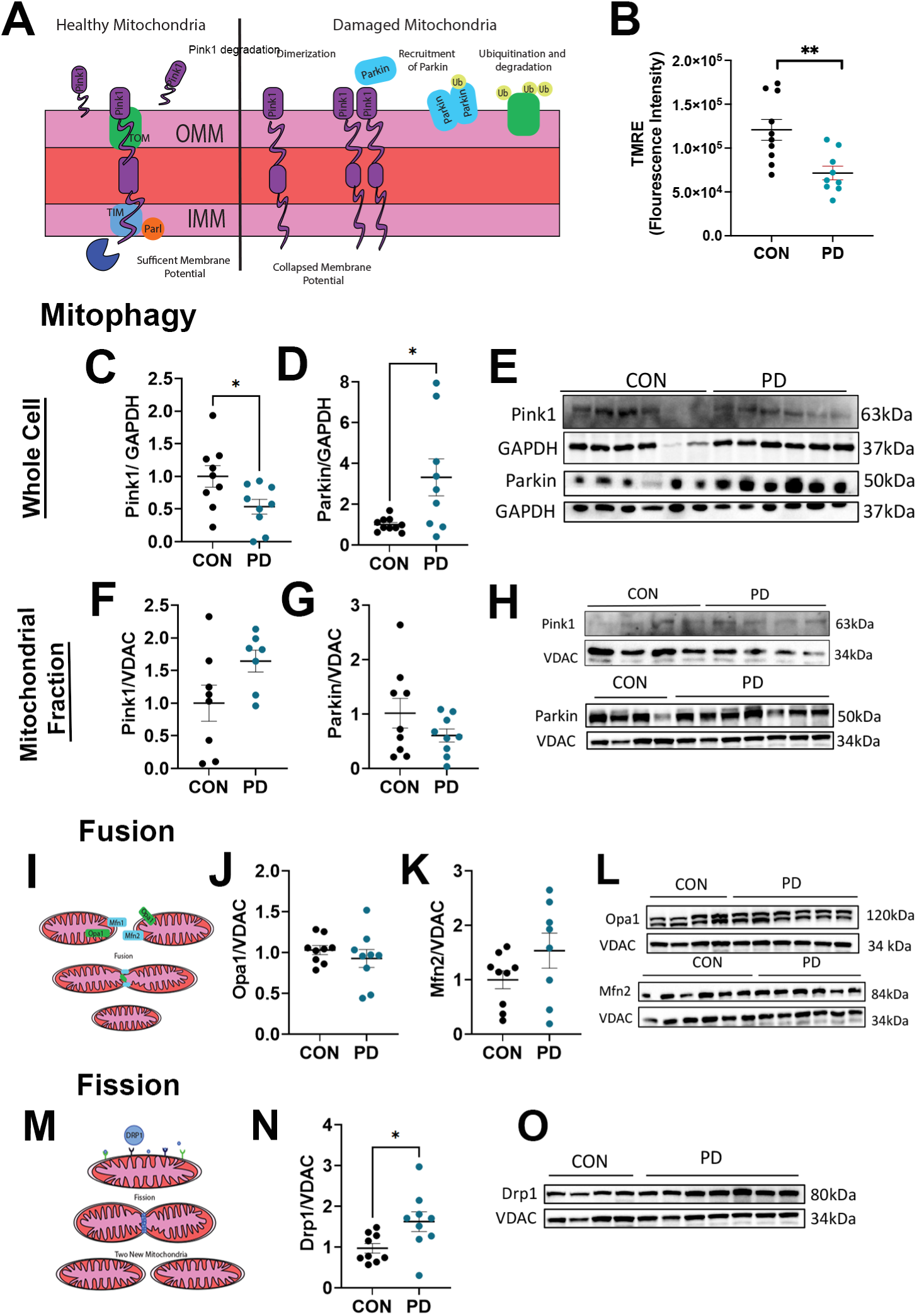
Mitochondrial dynamics in the iPSC-DAN. Schematic depiction of mitochondrial dynamics and degradation (A). Results from a TMRE assay elucidating changes in mitochondrial membrane potential are shown in (B). Whole cell lysates (C-E) and mitochondrial enriched fractions (F-H) from DAN were immunoblotted for mitophagy markers Pink1 (C,F; representative blots in E,H) and Parkin (D,G; representative blot in E,H). Cartoon of the process of Fusion is shown in (I). Fusion regulatory proteins Opa1 and Mfn2 were immunoblotted against and densitometrically quantified in (J-K). Representative blots are in (L). (M) shows a cartoon of the Fission process. Western blot results for Drp1, a protein involved in the control of fission, is shown in (N) with representative blots in (O). *P<0.05, **P<0.01, ***P<0.001, ****P<0.0001, Unpaired t-test; Mean ± SEM, N=8-10/group

We’ve previously shown the fragmentation of mitochondrial networks and altered morphology in PD iPSC-DAN (Corenblum *et al*., 2023). Mitochondria routinely undergo fission and fusion events in dynamic response to events within the cell, illustrated in **Fig. 5I, M** (Pernas and Scorrano, 2016; Tabara *et al*., 2025). Fusion events are regulated by the proteins mitofusin 2 (Mfn2), and optic atrophy-1 (Opa1). Fission, in which one mitochondrion is split into two is controlled by the protein dynamin-related protein 1 (Drp1). We probed the mitochondrially enriched fractions of iPSC-DAN and found no difference in the expression of fusion proteins between CON and PD cells (**Fig. 5J-L**). On the other hand, immuno-blotting of the fission protein, Drp-1, showed increased levels in the PD lines (1.624 ± 0.7244), compared to CON (0.970 ± 0.3472), (**Fig. 5N-O**; t= 2.443, df = 16, p = 0.0266, Unpaired t-test). These data suggested more exaggerated fission, and thus more fragmentation, in PD mitochondria.

### PD IPSC-DAN lines show accumulation of several forms of alpha synuclein

Aggregation and mis-localization of the synaptic protein alpha synuclein (αSyn) is a hallmark feature of clinical PD. We, and others have previously shown that reprogrammed iPSCs retain this feature of disease when differentiated into midbrain DA neurons (Corenblum *et al*., 2023; Oliveira *et al*., 2015; Zambon *et al*., 2019). Using antibodies targeted to monomeric αSyn by immunocytochemistry, we again show that PD iPSC-DAN have a greater expression of αSyn (**Fig. 6A-B**). Quantification of αSyn via NIH FIJI revealed that PD lines take up 7.22% ± 1.281 of each frame, while CON is 5.377% ± 0.952. (**Fig. 6C**; t= 3.462, df = 16, p = 0.0032, Unpaired t-test). Similarly, cell lysates were subjected to immuno-blotting for αSyn and phosphorylated αSyn (pSyn, pS129), (**Fig. 6D-G**). There substantially more αSyn (**Fig. 6D, F)** in PD lines (13.64±14.84, vs. 1.0± 0.8961; t = 2.7, df = 17, p = 0.0152, Unpaired t-test). There was also a greater amount of pSyn (**Fig. 6E-G**) in PD cells (20.57± 27, CON = 0.758 ± 0.611, t= 2.231, df= 16, p = 0.0332). In order, to get a better idea of the synuclein species within the cell, we also isolated Triton-X soluble, and Triton-X insoluble fractions of the cell (Anandhan *et al*., 2021). These fractions were then blotted directly on to nitrocellulose membrane and immuno-probed for both pSyn (**Fig. 6H**) and aggregated αSyn (**Fig. 6K**). These dot blots allowed for assessment of αSyn in its native form, without denaturing and linearization as required for electrophoresis. We saw a significant increase in pSyn in the soluble fraction (**Fig. 6I**) in the PD cells (3,389 ±551 a.u.) compared to CON (422,2±551 a.u.) (t = 5.535, df = 17, p < 0.001, Unpaired t-test). We saw a similar outcome within the soluble fraction for the aggregated αSyn (**Fig. 6L**), (CON = 818± 718.9 a.u. vs. PD = 2599± 1400 a.u.; t = 3.543, df = 17, p = 0.0025, Unpaired t-test). We saw no change to the total pSyn or aggregated αSyn in the insoluble fractions (**Fig. 6J, M**), indicating that these aberrant species are located mainly in the triton soluble fraction of cells. Taken together, these data point towards a dysregulation of αSyn within PD iPSC-DAN, and reminiscent of that found in disease, as indicated by the phosphorylated and aggregation specific detection.

**Figure 6:**
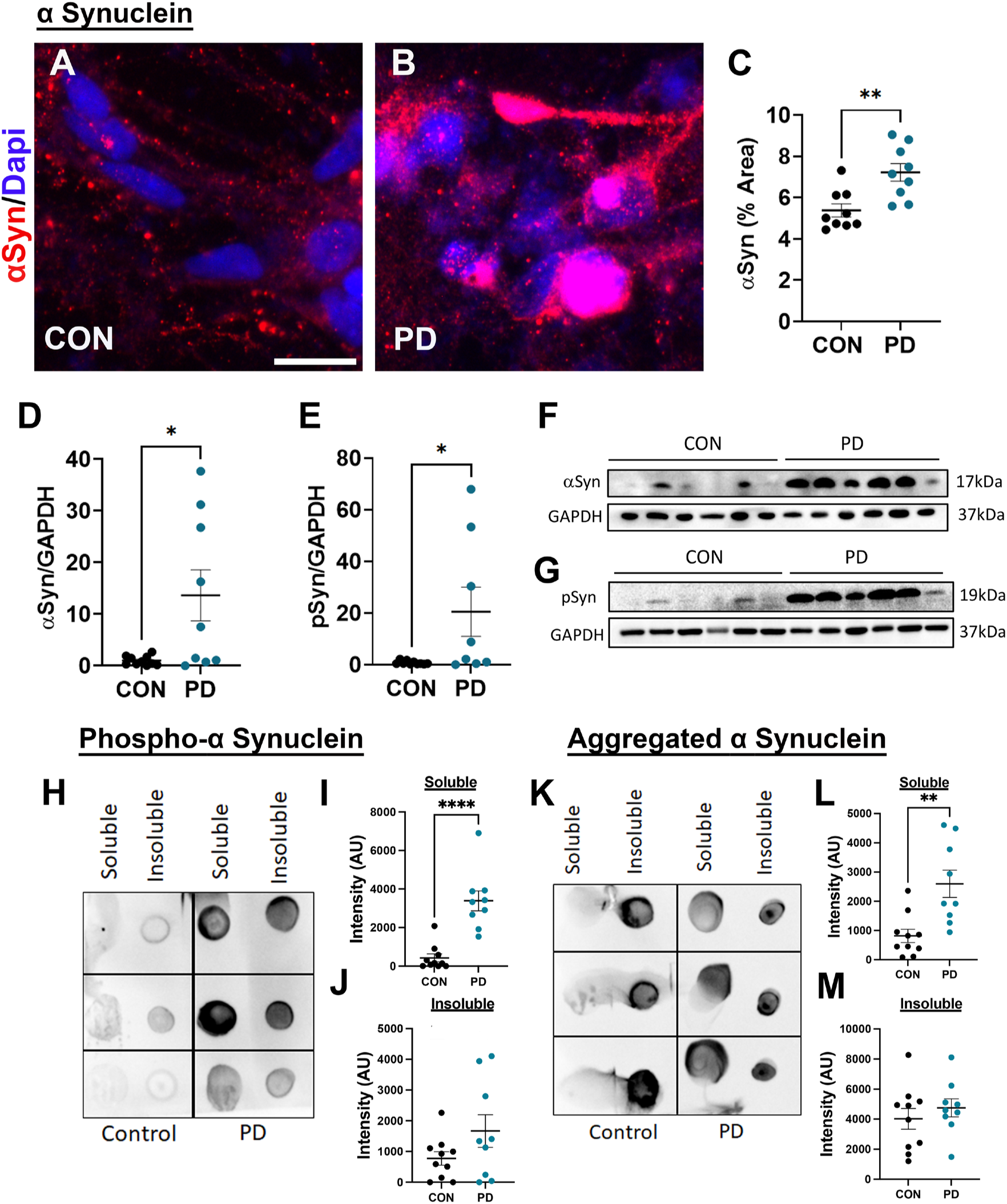
Alpha synuclein expression in the iPSC-DAN. Immunocytochemistry was used to assess monomeric αSyn in CON and PD DAN (A-B). Quantification of the percent area taken up by αSyn is shown in (C). Cell lysates were subjected to western blot analysis using αSyn and phosphorylated αSyn (pSyn) species (D-E). Representative blots are shown in (F,G). Soluble and Insoluble fractions were further blotted onto nitrocellulose directly and probed for both pSyn (H-J) and aggregated αSyn (K-M). Scale Bars: (A) = 10μm. *P<0.05, **P<0.01, ***P<0.001, ****P<0.0001, Unpaired t-test; Mean ± SEM, N=8-10 lines/group.

### Basal autophagy is increased in the PD DA neurons

Macroautophagy, a main form of autophagy, is an intercellular process of protein degradation (**Fig. 7A).** We have previously shown dysregulation of autophagic flux in idiopathic PD iPSC-DAN lines (Corenblum *et al*., 2023). In this study, using a separate and larger cohort of idiopathic PD iPSC-DAN, we assessed different proteins involved in autophagy. Under basal conditions, there were no changes between CON and PD lines for Atg4, a key early protein involved in lipidation of LC3II during phagosome formation (**Fig. 7B, D)**. However, there was an increase in the amount of total LC3II (**Fig. 7C, G**) in PD lines (CON = 1.0± 0.1998, vs. PD = 1.238± 0.1046; t = 3.045, df = 16, p = 0.0077, Unpaired t-test). P62 levels (**Fig. 7E, G**) did not show any significant alterations, and lysosomal marker, Lamp1 showed a trend towards reduced expression (**Fig. 7F-G**). These results suggest dysregulated autophagy in the PD iPSC-DAN.

**Figure 7:**
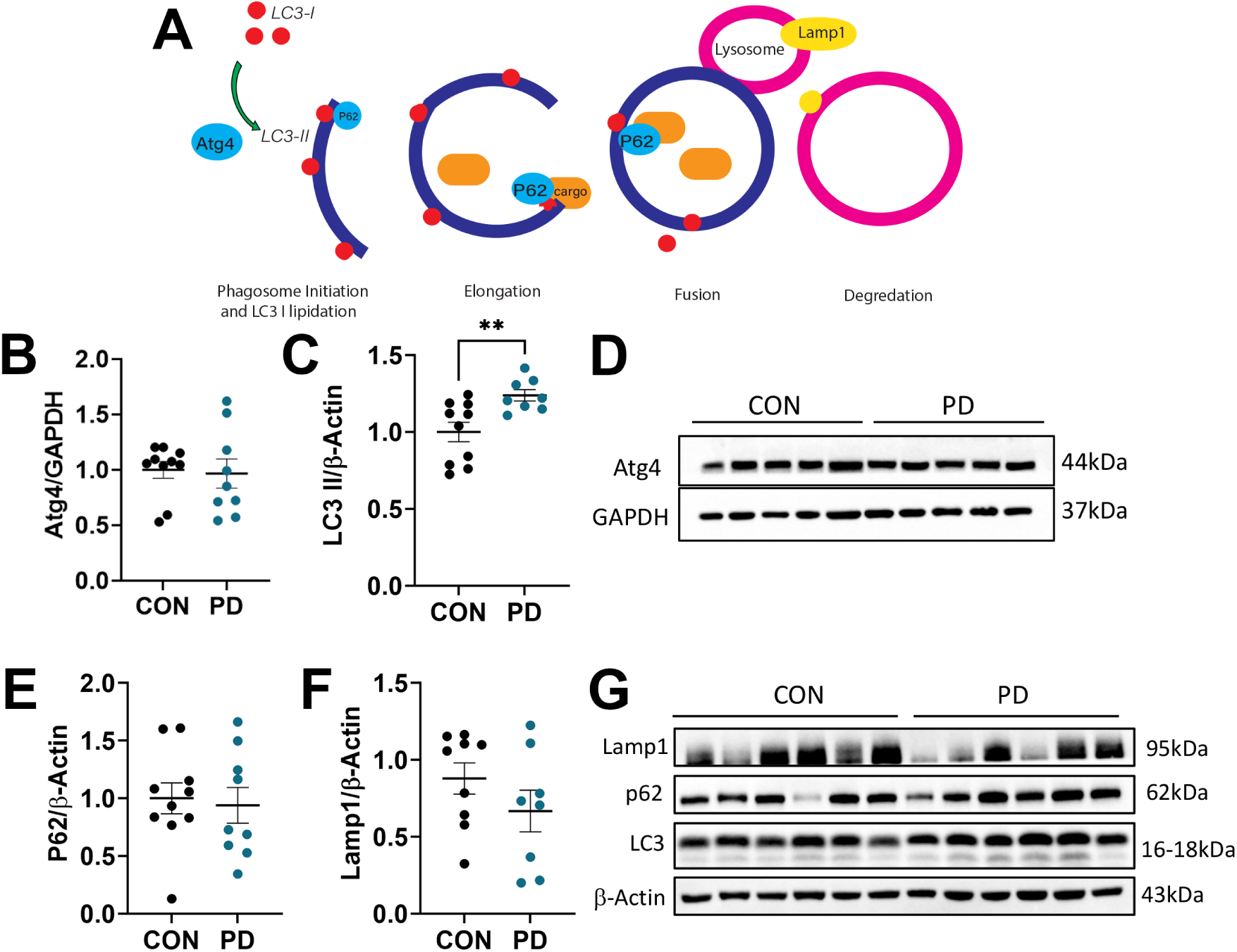
Changes in autophagy markers in the iPSC-DAN. (A) Schema illustrating the macroautophagy process. The levels of Atg4, a protein utilized in early lipidation during phagosome formation was quantified via western blot (B). Representative blots are shown in (D). Key autophagy associated protein LC3-II (C), adaptor molecule P62, (E) and autolysosome marker Lamp1 (F) were also examined. Example blots are depicted in (G). *P<0.05, **P<0.01, ***P<0.001, ****P<0.0001, Unpaired t-test; Mean ± SEM, N=8-10 lines/group.

### Alpha Synuclein directly associates with mitochondria in the PD iPSC-DAN

It has been proposed that αSyn can interact with mitochondria and lead to mitochondrial dysfunction (Di Maio *et al*., 2016; Geibl *et al*., 2024; Mingo *et al*., 2025). To assess the subcellular localization of αSyn, especially in the context of mitochondria and autophagic vesicles, we utilized immuno-electron microscopy. Day 52 DA neurons were immuno-labeled with antibodies for αSyn tagged to gold particles and subjected to transmission electron microscopy (**Fig. 8A-F,G-I**). There was a marked increase in the total area of αSyn gold particles associated with mitochondria in the PD iPSC-DAN (**Fig. 8A-B, G;** high mag images of mitochondria dotted with αSyn gold particles is in **C**) (t = 8.416, df = 6, p = 0.0002, Unpaired t-test). It was also seen that total area of αSyn associated with vesicles was significantly higher in the PD DA neurons vs CON (**Fig. 8D-E, H;** high magnification images of αSyn associated with vesicles is in **F**) (t = 4.349, df = 6, p = 0.0048, Unpaired t-test). Furthermore, the overall area of αSyn gold particles was higher in the PD DA neurons regardless of its subcellular localization, thus supporting the data in Figure 7 (**Fig. 8I**; t = 2.272, df = 6, p = 0.034, Unpaired t-test). Additionally, we used fluorescence microscopy to examine the association of αSyn with mitochondria via dual immunostaining for αSyn and TOM20, a marker of an outer mitochondrial membrane protein (**Fig. 8J-K**). There was no change to the total area of the TOM20-based mitochondrial network between CON and PD (**Fig. 8L**). However, we detected a strong increasing trend in percentage area of αSyn relative to the amount of positive mitochondria staining (**Fig. 8M**) (CON = 4.77 %± 3.765, PD = 8.632± 3.505, t =1.941, df = 14, p = 0.0726, Unpaired t-test). These data support the notion of increased αSyn-mitochondrial interactions and higher αSyn load in the PD iPSC-DAN.

**Figure 8:**
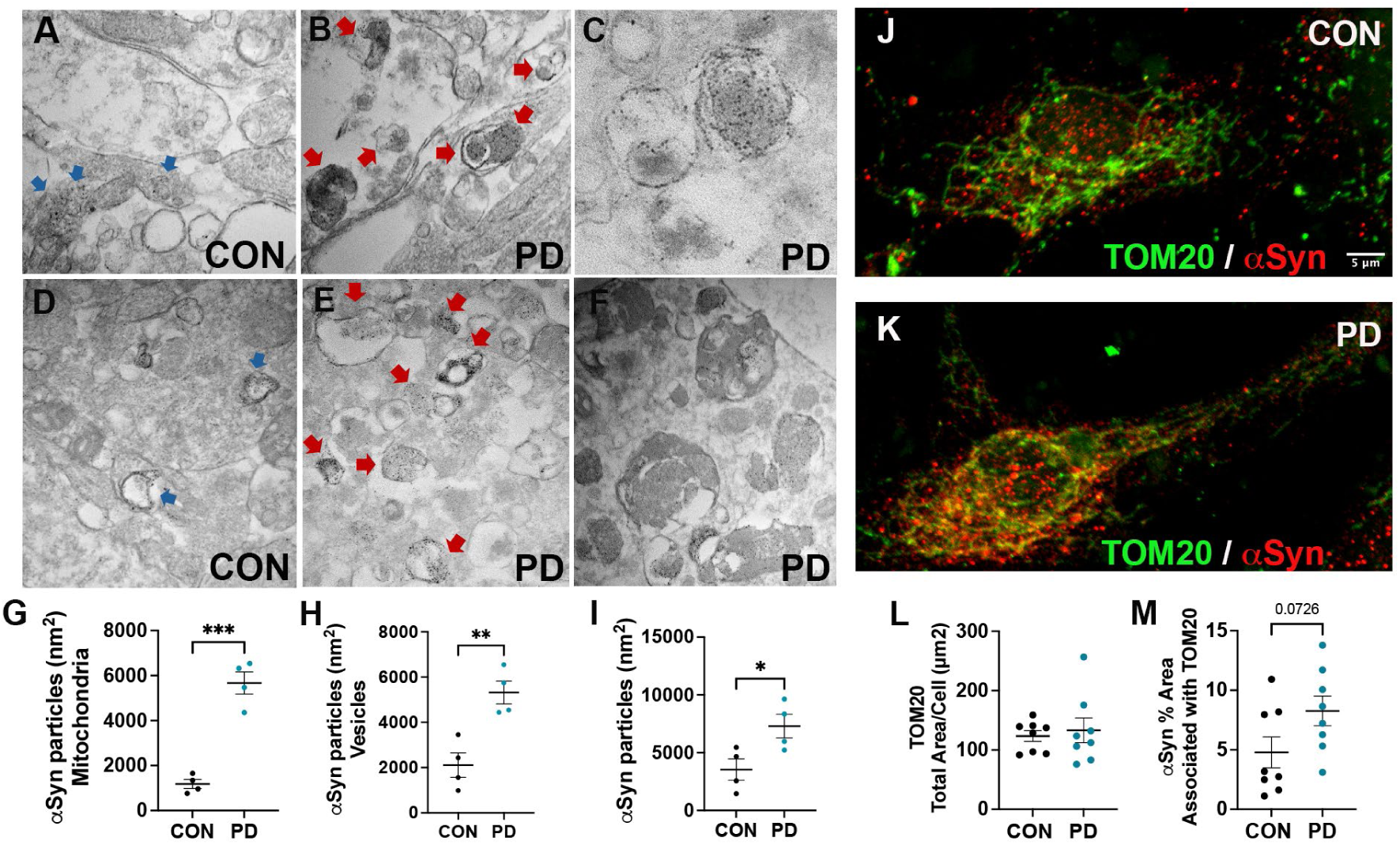
Subcellular localization of alpha synuclein in the iPSC-DAN. (A-F) Immuno-Electron Microscopy micrographs are shown for CON (A,D) and PD (B-C, E-F) DA neurons. The number of gold enhanced αSyn particles specifically associated with mitochondria are quantified in (G). Similarly, αSyn particles on vesicles is quantified in (H). The total amount of αSyn particles in the field shown in (I). Immunocytochemistry was utilized to assess colocalization of αSyn with mitochondria using TOM-20 (J-K). Total area of the TOM-20 mitochondrial network is quantified in (L). The amount of αSyn associated with positive TOM20 staining is shown in (M). Scale Bars: A-F = 250nm (scale bar for A,B,D,E is in D, scale bar for C & F is in F); J-K = 5μm and shown in J. *P<0.05, **P<0.01, ***P<0.001, ****P<0.0001, Unpaired t-test; Mean ± SEM, N=4-10/group.

## DISCUSSION

This study deeply analyzed bioenergetic and protein level alterations in human midbrain dopamine neurons derived from idiopathic PD subjects. We found that all the three major routes of ATP production and utilization, namely oxidative phosphorylation, glycolysis and creatine kinase, were compromised in the PD cells. This bioenergetic compromise also converged with altered mitochondrial membrane and fission/fusion dynamics, and impaired redox balance and protein quality control, ultimately affecting cellular survival. Fundamentally, these data unveil a network of mechanisms in human patient-derived DA neurons, that seemingly reinforce each other, contributing to neurodegeneration in the context of idiopathic PD.

Firstly, we detected across the board decreases in different aspects of oxidative phosphorylation function in the PD DA neurons in comparison with age and sex-matched controls. Application of the Seahorse extracellular flux assay showed that respiratory parameters such as the basal respiration, maximal respiration, proton leak, spare respiratory capacity, were all significantly reduced in PD DA neurons, thus reducing cellular ATP production. These findings aligned with our previous data, as well as data from others, that this vital metabolic process is affected in PD (Corenblum *et al*., 2023; Cunnane *et al*., 2020; Teves *et al*., 2018). When Glycolysis was probed, we did not see changes in rate of glycolysis or glycolytic capacity; however, the glycolytic reserve, which represents the cell’s potential to increase glycolytic activity in situations of increased demand, was notably lower in PD neurons. This implied that the PD cells were less able to upregulate glycolysis to meet increased ATP needs, when needed. Additionally, OCR to ECAR comparisons, showed that PD cells had lower OCR:ECAR ratios at basal and maximal respiration states, denoting a higher reliance on glycolysis than oxidative phosphorylation. Control cells on the other hand showed higher OCR:ECAR ratios. Additionally, the post-oligomycin ECAR and OCR:ECAR ratios (oligomycin inhibits mitochondrial respiration) displayed strong decreasing trends in the PD cells, suggesting that although glycolysis was less efficient in the PD neurons, the cells may be relying more on this process to produce ATP vs in control cells. Overall, these data indicate diminished oxidative phosphorylation and glycolytic capacities, with a somewhat greater dependence on glycolysis, in the idiopathic PD iPSC-DAN. Moreover, when creatine kinase B (CKB) levels were studied, lower amounts were noted in whole cell fractions, with higher amounts in the mitochondrial fractions of PD neurons. Increased CKB in the mitochondria indicates a potential disruption of energy metabolism - brain CKB is not typically found in high quantities in the mitochondria suggesting cell stress or malfunction where cellular compartmentalization of CKB is altered (Schlattner *et al*., 2016; Schlattner *et al*., 2006). Interestingly, the mitochondrial specific creatine kinase levels (uMt-CK) were not altered. These changes imply that the PD neurons may be compensating for their impaired energy state by increasing mitochondrial CKB. All in all, these data suggest weakened energy metabolism in the PD neurons at many scales.

Secondly, it was found that levels of Pink1 and Parkin, two mitophagy related proteins, were altered in the PD neurons. Mitophagy, is a selective form of autophagy that degrades damaged mitochondria (Pickrell and Youle, 2015). In fact, Pink1 and Parkin work together in a pathway that senses mitochondrial damage and initiates their removal through autophagy (Eiyama and Okamoto, 2015; Narendra and Youle, 2024; Pickrell and Youle, 2015). Under physiological conditions, Pink1 is imported to the mitochondria and quickly broken down, however, when mitochondrial membrane potential is lost, Pink1 accumulates on the outer mitochondrial membrane, and recruits Parkin, an E3 ubiquitin ligase (Narendra *et al*., 2008). The recruited Parkin can then ubiquitinate various mitochondrial proteins marking them for autophagic breakdown. Our data showed reduced mitochondrial membrane potential in the PD neurons paired with lower Pink1 in whole cell fractions and a strongly increasing Pink1 in the mitochondrial fraction, suggesting that Pink1 maybe accumulating to induce mitophagy of energetically compromised mitochondria in the PD neurons. Parkin on the other hand, was found to be increased in whole cell lysates but decreased in the mitochondrial fraction, signifying that although Pink1 may be collecting on the mitochondria it may not be recruiting Parkin efficiently. Overall, these data infer inefficient mitophagy and align with our previous findings in the same model, that showed greater mitophagic profiles and altered mitochondrial structure at the electron microscopic level in idiopathic PD neurons (Corenblum *et al*., 2023). Moreover, mitochondrial networks were also found to be fragmented in the Corenblum et al., 2023 paper. Supporting that concept, we found a higher expression of Drp1, a mitochondrial fission regulator, in PD neurons, supporting more mitochondrial fragmentation. On the other hand, the mitochondrial fusion proteins, Opa1 and Mfn2 were not different between PD and controls. Excessive fission/Drp1 activity in PD is well known and inhibiting this process has emerged as a therapeutic strategy (Feng *et al*., 2021; Filichia *et al*., 2016). It was shown that inhibition of Drp1 via a selective inhibitor, blocked its mitochondrial translocation and mitigated DA neuron loss in MPTP mice (Filichia *et al*., 2016). Feeding into all this, ROS levels were significantly increased, while the expression of GSTM-1, a key antioxidant enzyme, was significantly reduced in the PD neurons. GSTM-1 belongs to the class of glutathione-S-transferase (GST) enzymes, which are involved in detoxifying harmful substances, including oxidized dopamine (Vauzour *et al*., 2010). Specifically, GSTs reduce the harmful effects of electrophilic substrates through the transfer of glutathione. Thus, these data suggested that an imbalance between pro- and antioxidants was at play, thus enhancing oxidative stress in the idiopathic PD neurons. Nevertheless, it is unclear whether the ROS increase is due solely to the reduction of buffering capacity due to a decrease in GSTM-1, or simply due to greater ROS production from dysfunctional mitochondria. Interestingly, a meta-analysis has demonstrated that polymorphisms in GSTM-1 that result in a loss of function, increases susceptibility to PD (Weikang *et al*., 2016). Altogether, the results generate a picture of cascading events of oxidative stress, mitochondrial fragmentation and suboptimal mitophagy, causing insufficient mitochondrial energy production and the death of the PD neurons. Our data showing reduced TH expression (Supp figure 1) and viability (Fig. 1B) strengthen this notion.

Thirdly, the PD neurons displayed higher αSyn, and importantly, phospho-αSyn (pSerine129 epitope) than controls, as seen via immunostaining, immunoblotting and dot blots. Phosphorylation of αSyn is a significant event that is strongly associated with Lewy bodies and other pathological aggregates in PD (Barrenschee *et al*., 2017; Otto, 1969). This modification is thought to play a role in αSyn misfolding and aggregation (Awa *et al*., 2022). Thus, when we applied antibodies specific to aggregated αSyn, it was found that aggregated αSyn was indeed also notably elevated in the PD DA neurons. Along with these findings, on average, basal autophagy was found to be higher in PD cells (more LC3II). Interestingly, a strong trending towards lower LAMP1 expression was also seen, suggesting plausible lysosomal dysfunction, at least in some of the lines. Lysosomes are key to the macroautophagy process as they degrade the cellular materials in the autophagic vesicles. In fact, overwhelmed and inefficient autophagy, as well as lysosomal dysfunction, has been reported in PD (Fellner *et al*., 2021; Tang *et al*., 2021). Additionally, αSyn is a known target of autophagy and increased αSyn burden may in turn interfere with autophagy, reducing the degradation efficiency (Cuervo *et al*., 2004; Webb *et al*., 2003). αSyn aggregation can also negatively affect mitochondrial function and cell survival. To understand more, when ultrastructural quantification of αSyn gold particles associated with mitochondria and vesicles, via immuno-electron microscopy, was performed, we found significantly more such αSyn in PD DA neurons in comparison to controls (Fig. 8). αSyn was noted on mitochondrial and vesicular membranes as well as within the mitochondria and vesicles. These data implied direct interactions of αSyn with mitochondria and a greater vesicular load of αSyn in the PD cells. Moreover, on the mitochondrial front, confocal fluorescence analysis showed that there was more αSyn associated with TOM20, an outer membrane protein important in mitochondrial protein import, in the PD neurons. Previously, it has been shown that phosphorylated αSyn can bind to TOM20 and interfere with the import of mitochondrially targeted proteins (Di Maio *et al*., 2016). It is also known that Parkin binds to TOM20. Since we saw lower Parkin (in the mitochondrial fractions of the PD neurons, one possibility could be that the αSyn-TOM20 interaction was preventing Parkin from binding to TOM20 and affecting downstream mitophagy. Overall, our data suggests that more accumulated and aggregated αSyn may be directly impairing mitochondria, straining the autophagy system, and contributing to cellular stress and reduced viability of the PD DA neurons.

In summary, our study captures interconnected changes in different cellular energy systems, redox balance, and protein quality control mechanisms in human iPSC-derived midbrain dopamine neurons of idiopathic PD subjects. Malfunctioning of these pathways has been reported in PD animal models. However, investigations in human neurons are few, with most studies in iPSC-based neurons focused on genetic PD, which affects less than 10% of the PD population, rather than the more common idiopathic cases (>90%). Our work provides detailed information on these pathological pathways in idiopathic iPSC-DAN and how they may be linked. Given our previous proposition that iPSC-DAN models may convey early vulnerabilities in PD (Corenblum *et al*., 2023), this system offers a human platform to investigate early disease mechanisms in PD, which is an approach key to developing effective diagnostics and therapeutics.

## ACKNOWLEDGEMENTS

We thank Dr Moulon Luo for his technical assistance with the Seahorse mitochondrial assays and support from Dr Gunjan Goyal for the TMRE assay. The transmission electron microscopy was performed at the University of Arizona, ORP Imaging Cores - Electron facility, RRID: SCR_023279. The FEI Tecnai G2 Spirit BT TEM was supported by the grant from the NIH S10 OD011981. The cell lines used in this study were obtained from the Parkinson’s Progression Markers Initiative (PPMI) [www.ppmi-info.org]. PPMI, a public private partnership, is funded by the Michael J. Fox Foundation for Parkinson’s Research and corporate sponsors [https://www.ppmi-info.org/aboutppmi/who-we-are/study-sponsors]. The study was supported by a Michael J Fox Foundation Grant (MJFF 022138), and UA TRIF Core Pilot grant (ICEL-3771419) and intramural funds to LM.

## AUTHOR CONTRIBUTIONS

KB - Conception and design, Collection and assembly of data, Data analysis and Interpretation, Manuscript writing

MJC – Collection and assembly of data, Data analysis and Interpretation, Manuscript writing

PT – Conception and design (IEM), Collection and assembly of data (IEM), Manuscript writing

LM – Overall conception and design, Collection and assembly of data, Data analysis and Interpretation, Manuscript writing, Financial support, Final approval of manuscript.

## Data Availability

Data will be made available on request.

## SUPPLEMENTARY LEGENDS

**Figure Supp 1:**
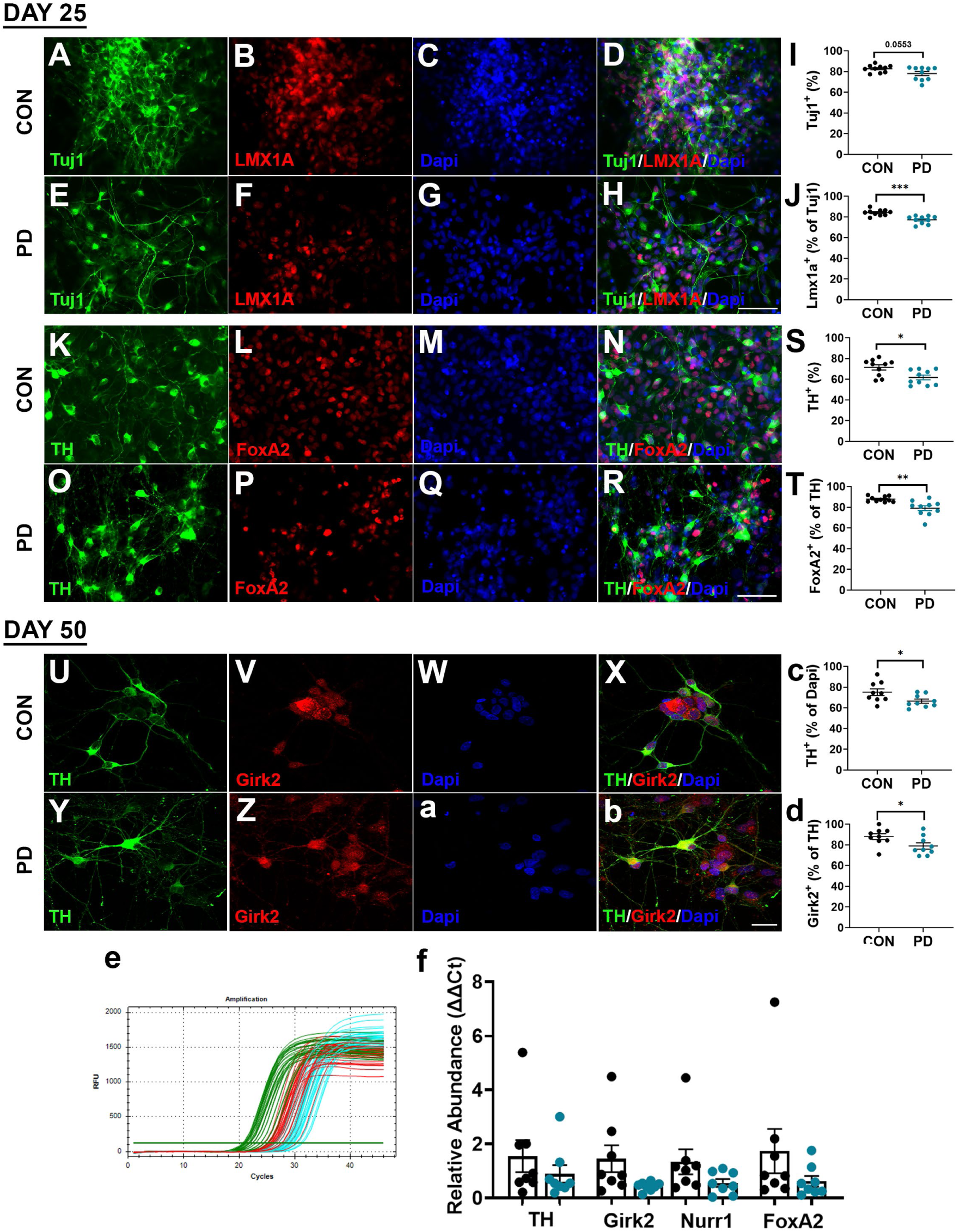
Characterization of iPSC-DAn lines from the patient fibroblasts at Days 25 and 50 of differentiation. iPSCs were differentiated (DA progenitor stage – Day 25) and matured into midbrain DA neurons (Day 50) using a modified dual-SMAD inhibition protocol. CON (A-D) and PD (E-H) cells were immunostained with the neuronal marker Tuj1 and midbrain DA neuron marker LMX1A, as well as neuronal marker TH and A9 subtype marker FoxA2 (K-N, O-R) at Day 25. Quantification of the efficiency of Tuj1^+^ and LMX1A^+^ is shown in (I,J). TH^+^ and FoxA2^+^ neuron production is shown in (S,T). After 50 days of differentiation, maturing neurons were immunostained and assessed for co-expression of TH/Girk2 to determine A9 subtype midbrain DA neuron generation efficiency (CON, U-X and PD, Y-b). Quantification is shown in (c-d). Example RT-qPCR expression curves are shown in (e). Enumeration of relative expression of maturing midbrain DA neuron markers TH, Girk2, Nurr1, and FoxA2 are in (f). Scale Bars: (A-H, K-R) = 50μm, (U-b) = 20μm. *P<0.05, **P<0.01, ***P<0.001, ****P<0.0001, Unpaired t-test, Mean ± SEM, N=8-10 lines/group.

**Figure Supp 2:**
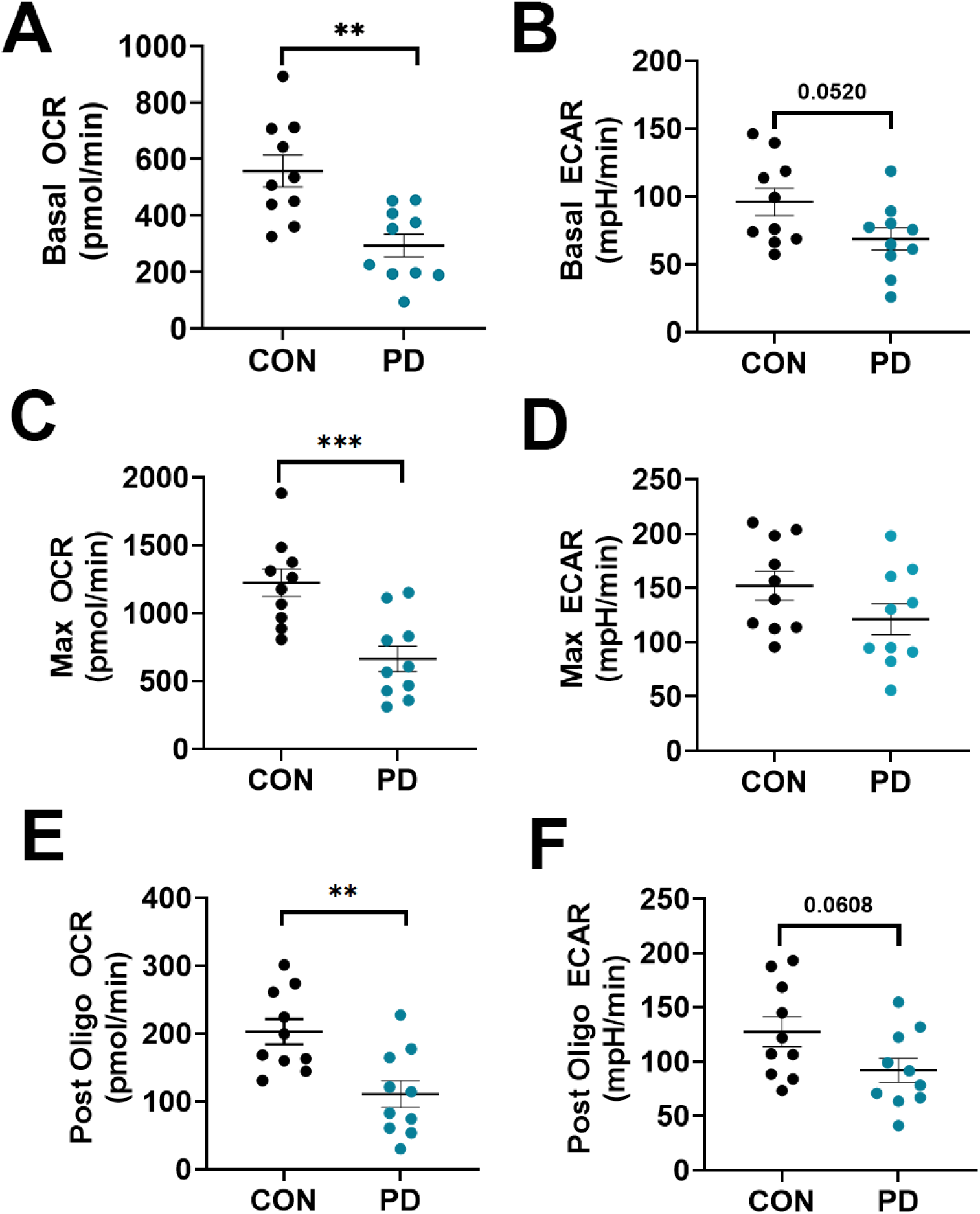
Raw OCR and ECAR values as proxy measurements to assess reliance on oxidative phosphorylation and glycolysis. Quantification of OCR and ECAR rates at baseline (A-B), maximal capacity (C-D) and post Oligomycin injection (E-F) are shown comparing CON to PD DA neurons. *P<0.05, **P<0.01, ***P<0.001, ****P<0.0001; Mean ± SEM, N=8-10 lines/group.

**Figure Supp 3:**
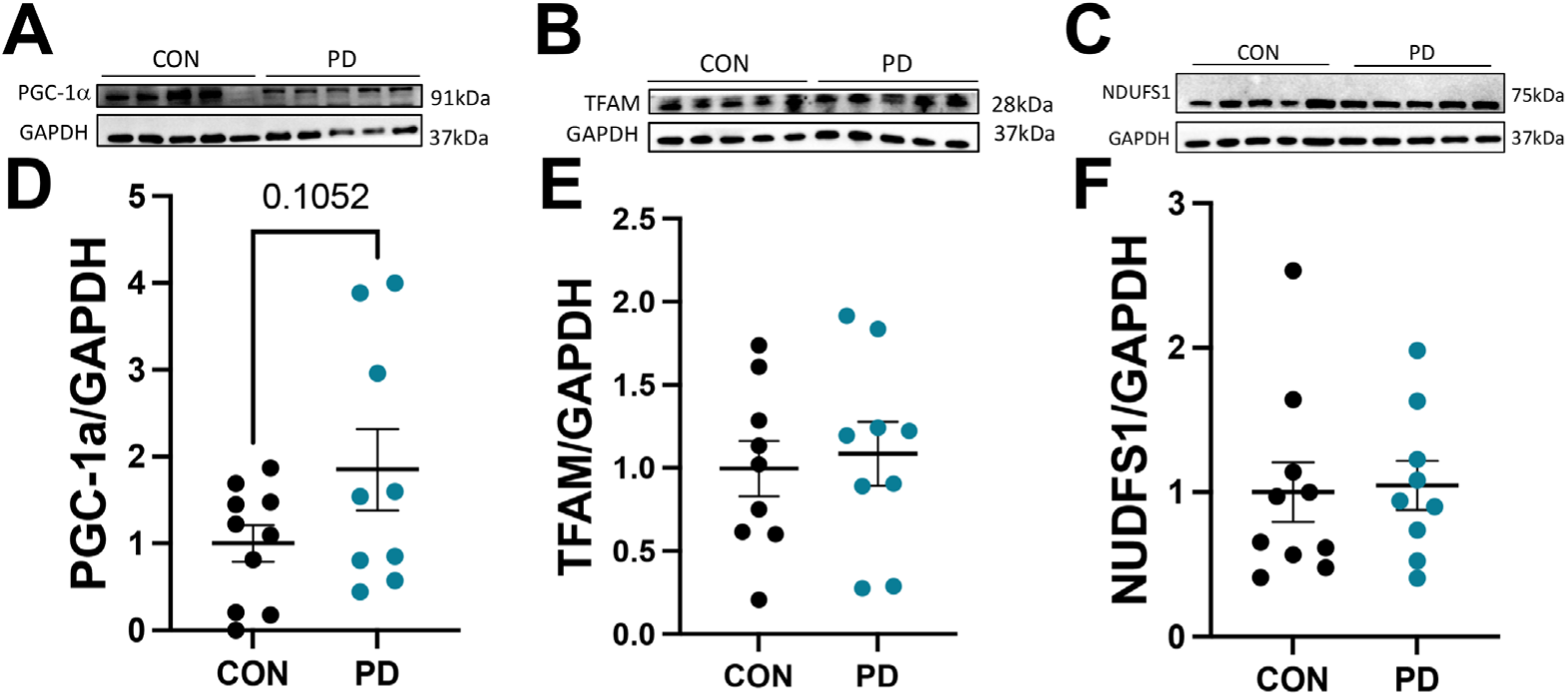
Mitochondrial biogenesis analysis. Western blot analysis was used to elucidate protein levels of mitochondrial biogenesis markers PGC1a (A), TFAM (B), and NDUFS1 (C). Quantification is in (D-F) respectively. *P<0.05, **P<0.01, ***P<0.001, ****P<0.0001, Unpaired t-test, Mean ± SEM, N=8-10 lines/group

## SUPPLEMENTARY TABLE LEGENDS

**Supplementary Table 1:** Line Description and Demographics. NA, not available; UPDRS, Unified Parkinson’s Disease Rating Scale Part 3 (on & off medications); UPDRS TOT, Unified Parkinson’s Disease Rating Scale Total (on & off medications);; NPI-Cog, Neuropsychiatric Inventory Score of Cognitive Impairment (0-Normal, 1-slight, 2-mild, 3-moderate, 4-severe); REM-Cat, REM Sleep Behavior Disorder Screening Questionnaire (RBDSQ) total score; A-Syn, CSF alpha-synuclein levels. Treatment is the participant on dopaminergic medication or receiving deep brain stimulation for treating the symptoms of PD. LED, calculated LED for each PD medication.

| Line ID | Group | Sex | Age at diagnosis (years) | Age at Blood Draw (years) | Disease Duration to Blood Draw (years) | UPDRS3 (ON) | UPDRS TOTAL (ON) | UPDRS3 | UPDRS TOTAL | NPI-Cog | REM | Treatment and LED (mgs) |
| --- | --- | --- | --- | --- | --- | --- | --- | --- | --- | --- | --- | --- |
| 3452 | CON | F | - | 71.3 | - | - | - | 4 | 8 | 0 | 4 | - |
| 3453 | CON | F | - | 67.8 | - | - | - | 2 | 5 | 0 | 3 | - |
| 3480 | CON | F | - | 76.3 | - | - | - | 2 | 15 | 0 | 3 | - |
| 3952 | CON | F | - | 74.4 | - | - | - | 0 | 5 | 0 | 3 | - |
| 3411 | CON | M | - | 45.6 | - | - | - | 1 | 5 | 0 | 4 | - |
| 3460 | CON | M | - | 75.6 | - | - | - | 6 | 11 | 1 | 2 | - |
| 3658 | CON | M | - | 62.6 | - | - | - | 0 | 0 | 0 | 6 | - |
| 3668 | CON | M | - | 79.0 | - | - | - | 0 | 5 | 0 | 8 | - |
| 3966 | CON | M | - | 63.4 | - | - | - | 0 | 1 | 0 | 5 | - |
| 4105 | CON | M | - | 69.8 | - | - | - | 0 | 1 | 0 | 1 | - |
|  | **Ratio F:M/ Mean** | **4:6** |  | **68.5** |  |  |  | **1.5** | **5.6** | **0.1** | **3.9** | **-** |
| 3419 | PD | F | 67.4 | 72.4 | 5.0 | 15 | 24 | 15 | 24 | 0 | 5 | NO |
| 3473 | PD | F | 55.4 | 59.4 | 4.0 | 46 | 94 | 50 | 98 | 2 | 7 | YES-1538 |
| 3664 | PD | F | 61.8 | 65.0 | 3.2 | 22 | 30 | 22 | 30 | 0 | 5 | NO |
| 3960 | PD | F | 52 | 56.8 | 4.8 | 9 | 22 | 19 | 32 | 0 | 1 | YES-598.75 |
| 4051 | PD | F | 52.9 | 56.1 | 3.2 | 26 | 35 | 26 | 35 | 0 | 3 | NO |
| 3234 | PD | M | 69.8 | 72.7 | 2.9 | 30 | 38 | 35 | 43 | 0 | 1 | YES-600 |
| 3459 | PD | M | 51.7 | 56.7 | 5.0 | 34 | 61 | 45 | 72 | 0 | 6 | YES-1000 |
| 3591 | PD | M | 62.7 | 66.7 | 4.0 | 19 | 29 | NA | NA | 0 | 3 | YES-960 |
| 3964 | PD | M | 39.6 | 44.6 | 5.0 | 6 | 18 | 7 | 19 | 0 | 3 | YES-360.1 |
| 4055 | PD | M | 69.9 | 72.9 | 3.0 | 33 | 56 | 33 | 56 | 0 | 5 | YES-100 |
|  | **Ratio F:M/ Mean** | **5:5** | **58.3** | **62.3** | **4.0** | **24** | **40.7** | **28** | **45.4** | **0.2** | **3.9** | **-** |

**Supplementary Table 2:** Detailed information for Immunoblots.

| Target Antigen | Protein Loaded (μg) | Gel Percent tage | Membrane | Company/Cat Number | Species | Dilution |
| --- | --- | --- | --- | --- | --- | --- |
| Whole Cell Lysate |  |  |  |  |  |  |
| aSyn | 12 | 12% | PVDF | BD-Biosciences #610786 | Mouse | 1:750 |
| pSyn | 17 | 12% | PVDF | Cell Signaling #23706 | Rabbit | 1:500 |
| Pink1 | 20 | 7.5% | NC | Biolegend #846201 | Mouse IgG1 | 1:1K |
| Parkin | 12 | 12% | PVDF | Santa Cruz SC-32282 | Mouse | 1:200 |
| LC3 | 8 | 12% | NC | Cell Signaling #4085 | Rabbit | 1:1K |
| Lamp1 | 8 | 12% | NC | Cell Signaling #3243 | Rabbit | 1:K |
| Atg4 | 8 | 12% | NC | Cell Signaling #13507 | Rabbit | 1:400 |
| P62 | 8 | 12% | NC | Novus H0000-878-M01 | Rabbit | 1:2K |
| GTSM | 15 | 10% | NC | Santa Cruz sc-55586 | Mouse | 1:200 |
| NQO-1 | 17 | 12% | NC | Santa Cruz sc-55586 | Mouse | 1:200 |
| HO-1 | 17 | 12% | NC | Santa Cruz sc-136960 | Mouse | 1:200 |
| γ-GCSm | 15 | 10% | NC | Santa Cruz sc-55586 | Mouse | 1:200 |
| CK-B | 10 | 12% | NC | Santa Cruz sc-373686 | Mouse | 1:200 |
| TFAM | 17 | 10% | NC | Abcam ab 131607 | Rabbit | 1:500 |
| PGC-1a | 15 | 10% | NC | Novus NPB1-04676PE | Rabbit | 1:500 |
| GAPDH | House Keeping |  |  | Cell Signaling #97166 | Rabbit | 1:2K |
| b-Actin | House Keeping |  |  | Santa Cruz sc-47778 | Mouse | 1:1K |
| Tubulin (DM1A) | House Keeping |  |  | Abcam ab 7291 | Mouse | 1:1K |
| Mitochondrial Fraction |  |  |  |  |  |  |
| Opa1 | 7 | 10% | NC | Cell Signaling #804715 | Rabbit | 1:1K |
| CK-B | 7 | 10% | NC | Santa Cruz sc-373686 | Mouse | 1:200 |
| Drp1 (D6C7) | 7 | 10% | NC | Cell Signaling | Rabbit | 1:1K |
|  |  |  |  | #8570 |  |  |
| Mfn2 (D2D10) | 7 | 10% | NC | Cell Signaling #9482 | Rabbit | 1:1K |
| uMt-CK | 7 | 10% | NC | Santa Cruz sc-514656 | Mouse | 1:200 |
| Parkin | 7 | 10% | NC | Santa Cruz SC-32282 | Mouse | 1:200 |
| Pink1 | 20 | 7.5% | NC | Biolegend #846201 | Mouse IgG1 | 1:1K |
| VDAC | House Keeping |  |  | Abcam Ab1-5895-1001 | Rabbit | 1:1K |
| Dot Blot |  |  |  |  |  |  |
| pSyn [MJF-R13 (8-8)] | 15 | NA | NC | Abcam Cat # Ab168381 | Rabbit | 1:500 |
| Aggregated aSyn [MJFR-14-6-4-2] | 15 | NA | NC | Abcam Cat # Ab209538 | Rabbit | 1:500 |
| Secondary Antibodies |  |  |  |  |  |  |
| HRP |  |  |  | Sigma #A0545 | Rabbit | 1:20K |
| HRP |  |  |  | Sigma #A9044 | Mouse | 1:20K |

**Supplementary Table 3:** Information on ICC Antibodies.

| **ANTIBODY** | **VENDOR** | **CATALOG #** | **HOST/TYPE/CLONE** | **CONCENTRATION** |
| --- | --- | --- | --- | --- |
| Anti-Tyrosine Hydroxylase | Millipore Sigma | MAB318 | Mouse/mAb/LNC1 | 1:2000 ICC |
| Anti-LMX1A | Millipore Sigma | AB10533 | Rabbit/pAb | 1:250 ICC |
| Anti-FoxA2/HNF3β | Cell Signaling Technology | 8186 | Rabbit/mAb/Gly138 | 1:200 ICC |
| Anti-Tuj1 | Millipore Sigma | MAB1637 | Mouse/mAb/TU-20 | 1:200 ICC |
| Anti-Girk2 (Kir3.2) | Alomone | APC-006 | Rabbit/pAb | 1:1000 ICC |
| Anti-Alpha Synuclein | BD-Biosciences | #610786 | Mouse/42 | 1:400 ICC |
| Anti-TOM20 (F-10) | Santa Cruz | sc-17764 | Mouse IgG2a | 1:400 ICC |

